# Population dynamics of generalist and specialist strategies under feast–famine cycles

**DOI:** 10.1101/2025.05.15.654191

**Authors:** Rintaro Niimi, Chikara Furusawa, Yusuke Himeoka

## Abstract

Microbial populations exhibit a broad spectrum of nutrient utilization strategies, ranging from those utilizing diverse nutrients, called “generalists,” to those being highly adapted to specific nutrients, called “specialists.” Identifying the conditions for the diversification of nutrient utilization strategies is one of the central questions in ecology. Previous theoretical studies have shown that trade-offs among different resource utilization functions in which cells cannot utilize broad types of substrates at nearly optimal efficiency are crucial for the emergence of diverse strategies. Additionally, in natural settings, nutrient availability often fluctuates over time, imposing another trade-off on the cells. Cells that grow rapidly under nutrient-rich conditions tend to have a higher death rate under nutrient-poor conditions, leading to a growth-death trade-off. This additional trade-off can contribute to the emergence of diverse strategies. Here, we introduce a mathematical model that simultaneously incorporates the resource-use trade-off and the growth-death trade-off. Nutrient supply was modeled as discrete stochastic events, mimicking temporal changes in nutrient availability. We show that the phenotype with a higher ratio of growth rate to death rate dominates the population; that is, the strength of the growth-death trade-off is a major determinant of whether generalists or specialists are dominant. We also found that a sparse and uncertain nutrient supply favors specialists, increasing their temporally averaged abundance. Our findings reveal that accounting for temporal environmental variation and the resulting growth-death trade-off is crucial for understanding the drivers of diversification in microbial nutrient utilization strategies.

**Author summary:** Microbial strategies for choosing the nutrients to utilize are highly diverse. In our study, we explored why some microbes become generalists able to use a variety of nutrients, whereas others specialize in only a few nutrients. These strategies often result from metabolic trade-offs during nutrient processing. However, in fluctuating environmental conditions, it is important to consider additional trade-offs; under nutrient-rich conditions, they can grow quickly, but this often comes at the cost of being more vulnerable when nutrients are scarce. To investigate how these trade-offs affect the emergence of generalists and specialists, we developed a simple model called the feast-famine cycle that mimics the natural cycles of plenty and scarcity of nutrients. Our work shows that the trade-off between growth and death rates is crucial for determining whether a generalist or specialist strategy is dominant. We highlight the importance of trade-offs in response to environmental changes in shaping microbial adaptive strategies.

## Introduction

Microbes are the most abundant and diverse organisms on Earth [1], inhabit a wide range of environments and form the fundamental basis of ecosystems [2]. Despite their diversity in the wild, the mechanisms that enable the stable coexistence of multiple microbial species remain an open question in mathematical ecology. Classical theories argue that increasing species diversity within an ecosystem tends to reduce stability, making the coexistence of species in a single habitat difficult [3, 4]. According to the classical interpretation of Gause’s competitive exclusion principle, species competing for the same resources cannot coexist and eventually lead to the dominance of a single species, driving others to extinction [5, 6]. Resolving such discrepancies between theories and observations has been a central question in the mathematical ecology field [7].

One key factor that facilitates species coexistence is the differentiation in nutrient utilization [7]. In the wild, nutrients for microbial cells are present in the form of complex molecules, not just as a specific carbon source such as glucose. These complex molecules may provide various carbon and amino acid sources. Species can avoid competition by utilizing different nutrients, enabling stable coexistence of multiple species within the same habitat [8, 9].

What drives the segregation of nutrient utilization by different microbial species? Why do no species monopolize all nutrients? Trade-offs play a pivotal role [10]. In laboratory experiments, microbes often exhibit trade-offs in their ability to utilize different nutrients, such as the trade-off between the utilization ability of fructose and galactose [11], silicon and phosphorus [12], and nitrogen and phosphorus [13]. When cells enhance their ability to utilize one nutrient, the utilization of other nutrients is constrained, which promotes differentiation of resource use among species. These trade-offs stem from various functional constraints on cellular metabolism, one of which is the limited capacity for protein abundance. A well-established view of proteome partitioning [14, 15] states that cells cannot activate all metabolic pathways at the maximum speed at the same time. The cells must decide which pathway to activate by sacrificing the capacity to utilize other nutrients [16]. Additionally, there are constraints due to the thermodynamic nature of metabolism. For instance, glycolysis and gluconeogenesis share several metabolic reactions, whereas the two metabolic pathways proceed in opposite direction. This makes the simultaneous utilization of glycolytic- and gluconeogenic carbons infeasible [11, 17].

The presence of such inherent trade-offs compels microbes to choose which nutrient to utilize, and may lead to a variety of nutrient utilization strategies. The two extremes of nutrient utilization strategies are *generalists* and *specialists*. Generalists can adapt to diverse conditions, but the utilization speed and/or efficiency of nutrients is typically lower than that of other strategies. Specialists use a narrow range of nutrients but achieve high growth rates in their adapted environments [18, 19]. Experimental and theoretical studies have provided several pieces of evidence that the emergence of diverse nutrient utilization strategies is instrumental in promoting ecosystem diversity [20–22].

Temporal variability of environmental conditions is another key aspect of the emergence of diverse ecosystems [23–27]. In natural environments, such as lakes, ocean surfaces, and soils, nutrients are not continuously supplied, as in laboratory chemostats, but rather appear in discrete and stochastic pulses. After nutrient input, the ecosystem undergoes a feast phase, which inevitably shifts to famine once nutrients are exhausted, a phenomenon known as the feast-famine cycle [28–31]. Moreover, not all nutrients are simultaneously available, and different nutrients often become available at different times. Therefore, cells must reorganize their proteome to exploit newly available resources, which requires the constitutive maintenance of additional nutrient-sensing and catabolic proteins. This incurs a growth cost [41, 42], and thus cells inevitably face an inherent resource-use trade-off in utilizing different nutrients.

Under feast-famine conditions, cells endure nutrient deprivation by reallocating resources from growth to stress protection. A key regulator is *rpoS*, which encodes *σ*^38^ (RpoS), the stress-response sigma factor in *E. coli*; deletion of *rpoS* (Δ*rpoS*) increases growth but compromises stress tolerance [33]. Because proteome remodeling upon sudden starvation is limited particularly in bacteria with relatively low protease activity [34], cells maintain basal expression of stress-response genes in advance, which ensures preparedness but reduces growth [37]. This allocation constraint leads to a trade-off between the growth rate during the feast period and the survival rate during the famine period. Indeed, a study by Biselli et al. [36] showed that there is a positive correlation between growth rate under nutrient-rich conditions and death rate in subsequent cultures without a carbon source.

How do these two trade-offs orchestrate and drive the diversity of nutrient utilization strategies? To date, the trade-off between the range of utilizable nutrients and the growth rate has been actively investigated and is considered one of the key factors of whether a generalist-like or a specialist-like strategy will have a higher advantage in a given environment; if strong trade-offs exist, where adaptation to one resource use significantly hampers the use of the other, specialists tend to evolve, and *vice versa* [16, 38, *39]*. Only with such a trade-off does the strength (i.e., the functional form of the trade-off), determine the fittest strategy. In this sense, environmental conditions, such as the amount of nutrients and frequency of supply, have no impact on the emergence of different nutrient utilization strategies.

However, the assumption of optimal strategies dictated solely by the functional form of trade-offs may be divorced from the ecological reality. Variations in nutrient availability and supply regimes strongly influence community composition and ecosystem processes in tropical ecosystems [40]. Therefore, we hypothesized that constraints arising from temporal variation in nutrient availability are key determinants of optimal nutrient utilization strategies. Henceforth, we examine the influence of two trade-offs—the trade-off between the range of utilizable nutrients and the growth rate, and the growth-death trade-off on the emergence of the multiple nutrient utilization strategy.

To determine how the resource-use trade-off and the growth-death trade-off synergistically affect nutrient utilization strategies, we modeled multiple phenotypes competing for nutrients under feast-famine cycles. In this model, nutrients are supplied by discrete and stochastic events that initiate the feast period. Once the feast period begins, cells consume nutrients to grow, leading to famine. We introduce two types of trade-offs for the microbes: if a phenotype utilizes a wide variety of nutrients, it cannot show growth as rapid as another phenotype that grows only on a limited number of nutrient sources. Additionally, if a phenotype has a higher growth rate during the feast period, it shows a higher death rate under famine conditions.

Using this model, we showed that either a generalist or specialist strategy can be dominant in the population, depending on environmental conditions. The transition of the fittest strategy from specialists to generalists occurred when the maximum average growth rate increased beyond the critical value, and the transition of the optimal strategy was triggered by the strength of the trade-offs. We also investigated the effect of fluctuations in nutrient supply intervals on nutrient utilization strategies. We found that specialists gain a relative advantage over generalists as supply becomes sparser and uncertain. Our results provide insights into a mathematical description of the diversification of resource use strategies in temporally varying environments.

## Model

In this study, we investigated the influence of trade-offs on the emergence of nutrient usage strategies in a fluctuating environment. To this end, we considered a population dynamics model for microbes in the feast-famine cycle. In the following, we use the term “phenotype” instead of species because different phenotypes differ only in their nutrient utilization strategy in the resolution of the present model.

The model consists of *N* phenotypes. Nutrients are supplied to the environment as a discrete stochastic event, whereas the population growth of the phenotypes is modeled using continuous-time deterministic ordinary differential equations. Here, we consider that there are multiple types of nutrients, including different carbon sources. There are *E* types of nutrient sources and a single type of nutrient is supplied by a single supply event. As will be seen, the supplied nutrient is almost always consumed much earlier than the next supply event, and thus, at most one type of nutrient is present in the system simultaneously. Hence, the type of nutrient supplied at the latest supply event sets the environmental condition; thus, we represent the environmental condition by the type of nutrient at the latest supply. Each nutrient supply event initiates a feast period during which cells consume the supplied nutrients and grow. Following nutrient depletion, the system transitions to a famine period, during which populations decline. We denote the growth rate and death rate of phenotype *i* in environment *e* by *µ*_*i,e*_ and *γ;*_*i,e*_, respectively. For a single environmental period, the model equation for the population of phenotype *i, X*_*i*_, is as follows:

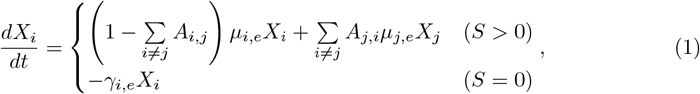

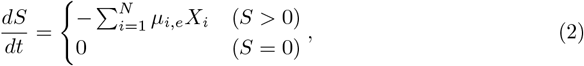

where *S* is the amount of nutrient. For simplicity, we assumed that the amount of nutrients was set to the type-independent value *S*_0_ at the supply event. As the type of supplied nutrient is encoded in the growth and death rate, we omit the subscript on *S* to denote the type of nutrient and only represent the amount. As noted earlier, the supplied nutrient is almost always exhausted well before the next pulse; accordingly, at most one nutrient type is present at any time, and no leftovers are carried over in our setup. The following results are not affected if we generalize the setup such that the nutrients can be left unconsumed. Because cells grow whenever nutrients remain, the population must experience the famine period for the average population level over a single feast-famine period. Thus, the supplied nutrients are almost always depleted before the next nutrient supply event after the population dynamics reach a steady state at the time-average level.

During the feast (*S >* 0) period, cells grow and proliferate at a rate *µ*_*i,e*_. During the division event of a cell of the *i*th phenotype, we allow the daughter cell to have the *j*th phenotype with a probability *A*_*i,j*_. During the famine (*S* = 0) period, cells die at rates *γ;*_*i,e*_. While we adopted a differential equation framework to describe continuous-time population dynamics, we incorporated the discreteness of the population by applying a threshold *θ* at each nutrient supply event. Any phenotype with a population size below threshold *θ* was set to zero. This threshold was evaluated only at nutrient supply events. Note that *X*_*i*_ = 0 is not the absorbing state for the *i*th phenotype because the population is supplied from other phenotypes by the phenotypic switching process.

Nutrient supply events occur at random and discrete time points *τ*_*k*_ (*k* = 1, 2, …). At each event, the amount of nutrient *S* discontinuously jumped to a constant value *S*_0_.

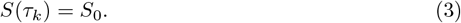

The type of supplied nutrient, that is, the environmental index *e*, was randomly chosen for the nutrient supply event *τ*_*k*_ (Fig 1).

**Fig 1.**
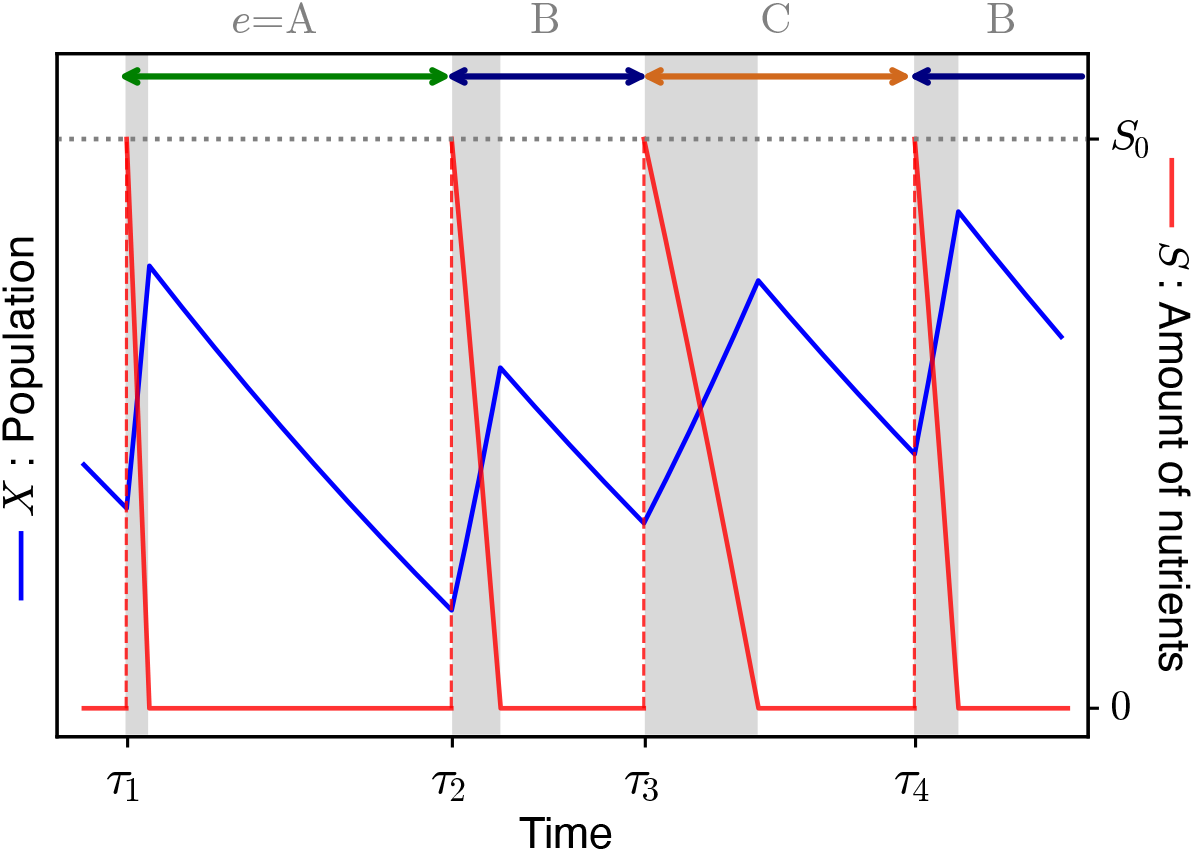
Schematic of the population dynamics model. Temporal changes in population size and nutrient levels in a single phenotype (*N* = 1) case. The blue line represents the population *X*, and the red dashed line represents the amount of nutrients *S*. At the time points *τ*_1_, *τ*_2_, ·, nutrients are supplied and *S* is set to *S*_0_. The feast phase (gray shaded area) begins at each nutrient supply event and ends when the supplied nutrient is exhausted by cellular consumption, after which the system enters the famine phase. At each time *τ*_*k*_, a different type of nutrition is randomly selected and supplied. The type of nutrients supplied depends on the value of environmental variables. Here, we have only a single phenotype and the growth rates are set such that *µ*_*A*_ *> µ*_*B*_ *> µ*_*C*_.

In this model, we introduced two types of trade-off. The first trade-off is the resource use trade-off.

Here, we assume that there is a strong trade-off in the cost of utilizing different nutrients. When two distinct cellular traits *x* and *y* have a trade-off relation *y* = *f* (*x*), the sign of the second derivative of the trade-off function (*d*^2^*y/dx*^2^) is often used as a measure of the strength of the trade-off [38, 39]. When the trade-off function is convex (*d*^2^*y/dx*^2^ *>* 0), the acquisition of both traits is more restricted than in the marginal, linear trade-off case (*d*^2^*y/dx*^2^ = 0). Therefore, this is referred to as a strong trade-off. However, when the trade-off function is concave (*d*^2^*y/dx*^2^ *<* 0), it is called a weak trade-off. According to Caetano et al. [16], there is no room for nutrient utilization strategies to diversify with weak trade-offs [16]. Thus, we introduced a strong trade-off between the growth rates of different nutrients using the following function, while the specific choice of the function did not alter the result (see S1 Text for details).

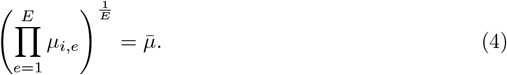

In addition to the above trade-off (Eq (4)), we introduce a trade-off between growth and death rates. In a study by Biselli et al. [36], the authors cultured *E. coli* cells in several flasks containing distinct sole carbon sources, and then the cells were washed and transferred into a carbon-free buffer medium to examine the relationship between growth rate under nutrient-rich conditions and death rate during the subsequent starvation state. They reported that the death rate in a carbon-free buffer medium increases exponentially with the growth rate under nutrient-rich preculture conditions. This relationship was generic for the different carbon sources in the medium. According to this result, we introduce an exponential trade-off between the growth rate and the death rate as follows:

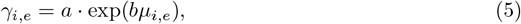

where *a* and *b* are constant values independent of the indices of phenotype (*i*) and environment (*e*), respectively.

## Results

### Transition between Generalist-dominance and Specialist-dominance

The purpose of this study was to identify the factors that drive the diversification of nutrient utilization strategies in temporally varying environments. As an initial step for answering this question, we start with a simple setup, where the model equation consists of *N* = 3 phenotypes and *E* = 2 environmental conditions. We index the phenotypes by Arabic numbers (phenotypes 1, 2, and 3) and environments by alphabet (environments A and B).

In this simple setup, we set the growth rates of phenotypes in ascending and descending order, that is, the growth rate in environment A follows *µ*_1,*A*_ *> µ*_2,*A*_ *> µ*_3,*A*_ and that in environment B follows *µ*_1,*B*_ *< µ*_2,*B*_ *< µ*_3,*B*_. To begin with a simple case, we assume symmetry between environments A and B by max_*i*_ *µ*_*i,A*_ = max_*i*_ *µ*_*i,B*_ and min_*i*_ *µ*_*i,A*_ = min_*i*_ *µ*_*i,B*_. One way to implement this symmetry consistent with the resource-use trade-off (Eq (4)) is as follows:

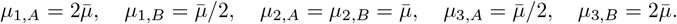

Because of this growth rate setting, phenotype 2 has a generalist-like strategy, where the growth rates are the same in the two environments, whereas the other phenotypes are specialist-like compared to phenotype 2 (see Fig 2A). Note that the two specialists can also grow under environmental conditions in which the phenotypes have a lower growth rate, therefore, they are not extreme specialists who cannot grow under certain environmental conditions.

**Fig 2.**
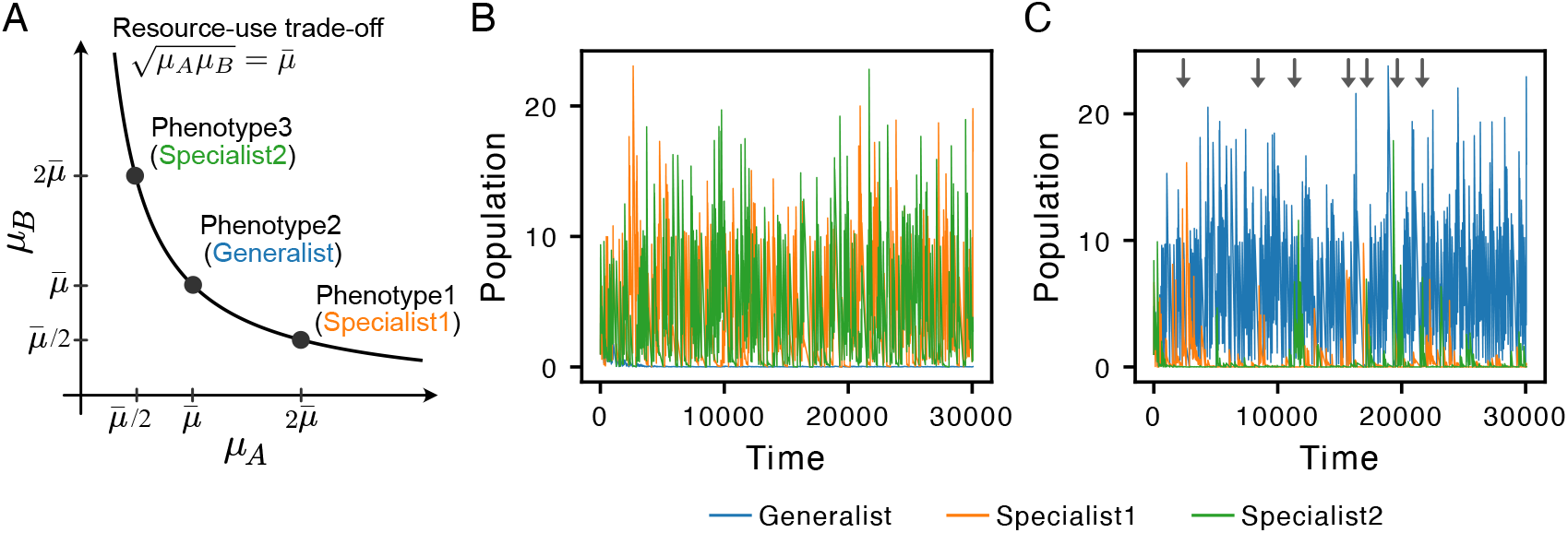
Examples of the simulation with *N* = 3 and *E* = 2. (A) The generalist (phenotype 2) is at 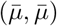; the two specialists (phenotypes 1 and 3) are symmetric at 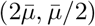 and [ineq} respectively. (B) The case with 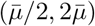. Two specialists dominate the population. (C) The case with 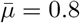. The generalist dominates the population. The initial values of populations are set to unity for all three phenotypes. The parameters are set to *a* = 0.01, *b* = 1, *S*_0_ = 10, *p* = 10^−4^, and *θ* = 10^−8^. The waiting time for the nutrient supply Δ*τ*_*k*_ (= *τ*_*k*+1_ − *τ*_*k*_) follows the gamma distribution with shape and rate parameters being 2 and 50, Δ*τ*_*k*_ ∼ Γ(2, 50).

We assumed that the choice of the nutrient utilization strategy did not affect the probability of phenotypic switching. In addition, the phenotypic switch is considered to occur only in the nearest neighbor of the phenotype index. Overall, the phenotype switching probabilities are given by

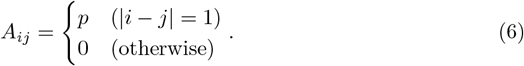

We set the waiting time for nutrient supply events, Δ*τ*_*k*_ to follow a gamma distribution. The gamma distribution is a generalized form of the Erlang distribution that is widely employed to describe the queuing time distribution of events that follow the Poisson process.

Fig 2 shows the population dynamics in two different simulation setups: 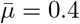 and 0.8. We observed two distinct types of population dynamics: in the first case, two specialist phenotypes dominated the community (Fig 2B), and in the second case, the generalist phenotype became dominant (Fig 2C). This result was also observed when nutrient supply events occurred at regular intervals Δ*τ*_*k*_ = Δ*τ* = const. (S1 Fig).

To understand how the dominant phenotypes changed depending on the geometric mean of the growth rate 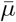, we simulated the model with various values of 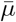. In this simulation, we set the intervals between successive nutrient supply events Δ*τ*_*k*_ constant to clarify the effect of the parameter value on population dynamics. Fig 3A shows the temporal average of the population of the generalist and two specialists. The temporal average of the two specialists was taken, and the mean of the two averages is plotted in the figure. The generalist is dominant for a larger 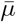, and the specialists are dominant for a smaller 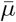.

**Fig 3.**
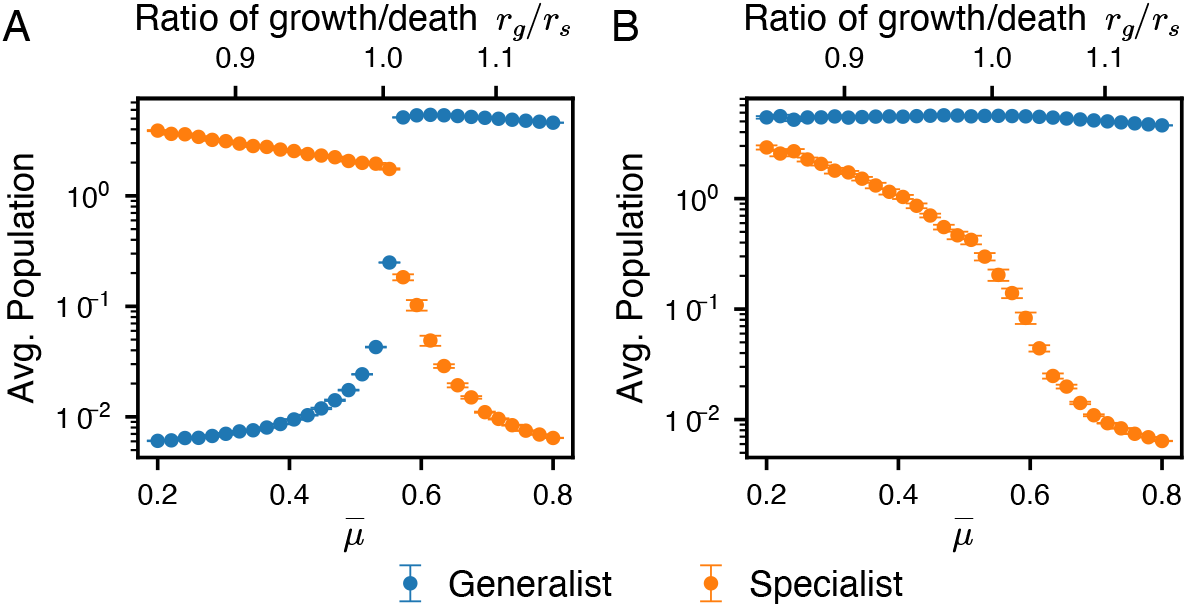
Temporal average of the population with constant Δ*τ*. The temporally averaged population are shown as a function of the parameter 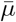. The model with one generalist and two specialists is simulated for (A), while there is only a single specialist adapted to environment A for (B). We simulated up to *t* = 3.0 10^5^ and averaged the populations after *t* = 2.5 10^5^. For each 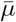, we ran 10 simulations and averaged the results. The error bars indicate the standard error. The secondary *x*-axis shows the ratio of *r*_*g*_ to *r*_*s*_. Throughout the simulations, the strength of the growth-death trade-off was fixed at *b* = 1, and nutrient supply events occurred at constant intervals (Δ*τ* = 100). All other parameters were identical to those used in Fig 2.

To elucidate the mechanism of the transition between generalist and specialist dominance, we introduce a simple dimensionless parameter given by *r* = ⟨*µ*⟩*/* ⟨*γ*⟩, where ⟨*µ*⟩ and ⟨*γ*⟩ are given by ⟨*µ*⟩= (*µ*_*A*_ + *µ*_*B*_)*/*2 and ⟨*γ*⟩ = (*γ*_*A*_ + *γ*_*B*_)*/*2, respectively. Because the two specialist phenotypes exhibited symmetric growth properties with respect to the two environments (*µ*_1,*A*_ = *µ*_3,*B*_ and *µ*_1,*B*_ = *µ*_3,*A*_), the values of *r* for the two specialists were the same. We denote the *r* values of the specialists and generalists as *r*_*s*_ and *r*_*g*_, respectively. The ratio of *r*_*g*_ to *r*_*s*_ was used as the horizontal top axis in Fig 3A. The figure shows that the transition between generalist- and specialist-dominant dynamics occurs at the value of *r*_*g*_*/r*_*s*_ being unity. The value of *r*_*g*_*/r*_*s*_ at the transition point is robust to the parameter values of the phenotype switching probability *p* (S3 Fig). The transition of dominant phenotypes was well captured by the ratio of the *r*-value between generalists and specialists. We carried out the same analysis by simulating the population dynamics by changing *b* value, which represents the strength of the growth-death trade-off (S2 FigA).

To determine why the ratio *r*_*g*_*/r*_*s*_ determines the dominant phenotypes, we analytically estimated the transition point (see S1 Text for details). Here, we examine the stability of generalists or specialists by calculating the invasibility of the other phenotypes. We examined the invasibility of an invasive phenotype introduced at low frequency. We demonstrated that the invasive phenotype was fixed when the ratio of the growth rate to the death rate was larger than that of the preexisting phenotype.

Given that *r* is defined as the ratio of the arithmetic mean of growth rate across the two environments to the arithmetic mean of death rate across the two environments, we therefore perform our analysis in a single, “averaged” environment by substituting *µ* and *γ* with their respective arithmetic means across the two environments. Phenotype *i* has a single average growth rate *µ*_*i*_ and death rate *γ*_*i*_ defined as follows:

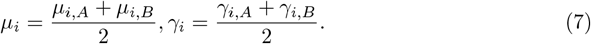

To determine the capacity of each phenotype to persist and invade under time-varying resource availability, we summarized the overall population change across a single feast-famine cycle by net log growth between two consecutive nutrient supply events.

The net log growth of phenotype *i, f*_*i*_ is given by

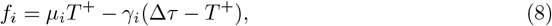

where *T* ^+^ is the duration of nutrient availability after the nutrient supply. If *f*_*i*_ is positive, the population size increases, and if *f*_*i*_ is negative, the population size decreases. When there is only a single phenotype (phenotype *α*), the population size settles at the point where *f*_*α*_ = 0. Here, we considered the invasibility of an other phenotype (phenotype *β*). Phenotype *β* can be fixed when *f*_*β*_ is positive. Because phenotype *β* is a mutant phenotype introduced at a low frequency, the population of phenotype *β* is sufficiently small, and thus, the change in the value of *T* ^+^ is negligible. Because the value of *T* ^+^ satisfies *f*_*α*_ = 0,

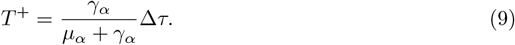

Thus, *f*_*β*_ is given by

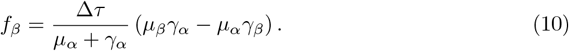

Therefore, phenotype *β* can invade if *µ*_*β*_*/γ*_*β*_ > *µ*_*α*_*/γ*_*α*_ holds.

Interestingly, the transition did not occur in a simulation with one type of specialist and one type of generalist (*N* = 2, *E* = 2, phenotype 3 excluded) (Fig 3B and S2 Fig B). In such cases, a generalist always dominates the system, and thus, the transition from generalist dominance to specialist dominance takes place through the cooperation of specialists to suppress the increase in the generalist population.

In the present simulation, we studied a simple model with one generalist and two specialist phenotypes. Phenotype switching doubles the growth rate in one environment and halves it in the other. Switching occurred between the three discrete phenotypes. However, in natural evolutionary processes, the effects of phenotypic change are typically gradual and continuous. To ensure that our findings are not an artifact of the limited number of phenotypes, we repeated the simulations with a finer discretization of phenotypes, where growth rates vary smoothly along the constraint surface 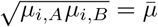(see the last part of the Results section). This refined setting yielded a similar outcome: the system still underwent shifts in dominance between the generalist and specialist strategies. Hence, the conclusions reported here are robust to the granularity of phenotypic space.

### Specialists have advantage under sparse and uncertain nutrient supply

#### Mean of nutrient supply interval

So far, we have seen the effect of the two parameters 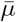 and *b* on population dynamics, which determine the ratio of the arithmetic mean of the growth rate to that of the death rate for the generalist and the specialists, *r*_*g*_ and *r*_*s*_. In this section, we investigate the effect of the nutrient supply interval Δ*τ* on population dynamics.

Fig 4 shows the temporal average of the population of the generalist and two specialists as a function of Δ*τ* . The effect of Δ*τ* on the temporally averaged population depends on whether the system is in the specialist-dominant or generalist-dominant condition, which is determined by the value of 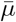. In the case of 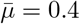 (Fig 4A, specialist-dominant condition), the temporally averaged population of the generalist and the specialists declined as Δ*τ* increased. This decline is expected because each supply event delivers a fixed amount of nutrient *S*_0_; therefore the time-averaged nutrient supply rate over a long window is *S*_0_*/*Δ*τ* . As Δ*τ* increases, the time-averaged nutrient supply rate decreases, which lowers the temporally averaged population.

**Fig 4.**
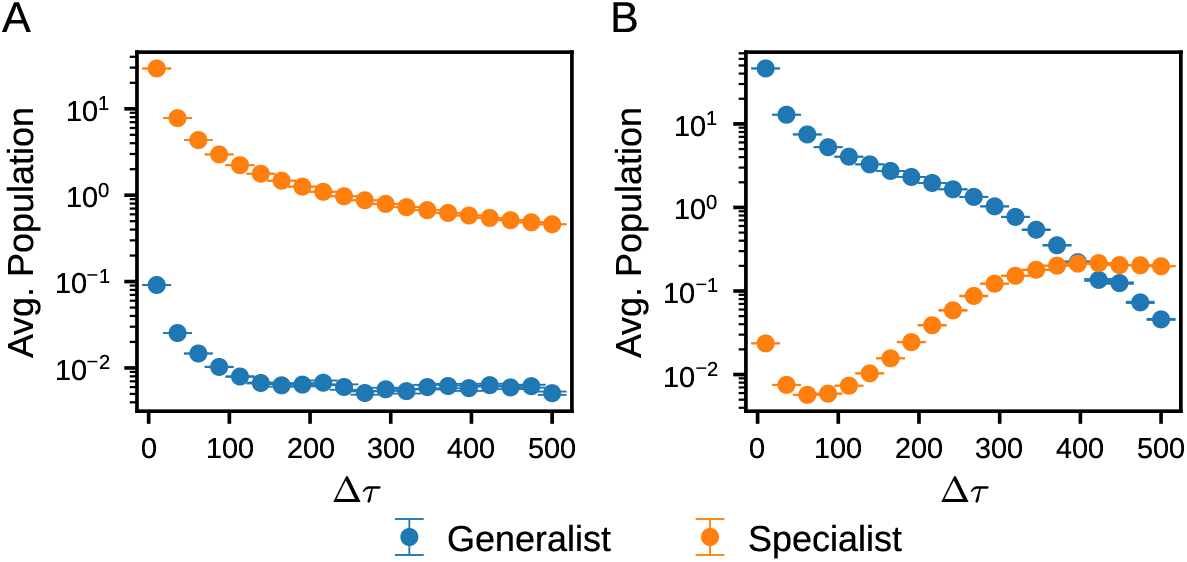
Temporal average of the population as a function of Δ*τ*. Changes in the temporally averaged populations as the parameter Δ*τ* is varied in simulation with one generalist and two specialists. The case with 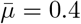 is shown in (A) and the case with 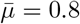 is shown in (B). We simulated up to *t* = 3000 × Δ*τ* and averaged the populations after *t* = 2500 × Δ*τ* . Δ*τ* is constant value. For each Δ*τ*, we ran 10 simulations and averaged the results. The error bars indicate the standard error. All other parameters were identical to those used in Fig. 2.

On the other hand, in the case of 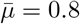 (Fig 4B, generalist-dominant condition), the temporally averaged population of the specialists increased within a certain range of Δ*τ*, while that of the generalist continued to decline. For a sufficiently large Δ*τ*, the temporally averaged population of the specialists exceeded that of the generalist.

To explain the counterintuitive increase in specialist population under generalist-dominant conditions, we analytically estimated the temporally averaged population and quantified the effect of Δ*τ* on the temporally averaged population. To facilitate the analysis, we assumed that the specialist population was sufficiently small that phenotypic changes from specialists to generalists were negligible. There were three phenotypes (phenotype 1,2 and 3) and two environmental conditions (environment A and B). Phenotype 1 was a specialist in environment A; phenotype 3 was a specialist in environment B, and phenotype 2 was a generalist. We assumed that nutrient supply events occurred at constant intervals Δ*τ* and that the environment *e* changed alternately. Under this deterministic condition, the population dynamics asymptotically converge to a periodic orbit. We analytically computed the temporally averaged population size of the periodic solution. The detailed derivations and results are provided in S1 Text.

From our calculation, we roughly estimated the order of change in the population of generalists and specialists with respect to Δ*τ* (see S1 Text for details). The temporally averaged populations have different dependencies on Δ*τ* in the small-Δ*τ* and large-Δ*τ* regimes, as follows:

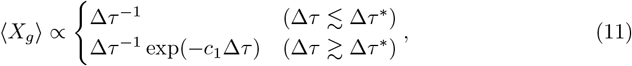

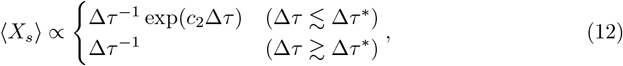

where *c*_1_ and *c*_2_ are positive constants on the order 10^−2^. Δ*τ* ^∗^ is the characteristic timescale separating the two regimes.

Both generalist and specialist populations exhibit a Δ*τ* ^−1^ decline because the nutrient influx per unit time likewise decreases as Δ*τ* ^−1^. According to Eq (12), when Δ*τ* ≲ Δ*τ* ^∗^ holds, the temporally averaged population of the specialists is amplified by an exponential factor, exp(*c*_2_Δ*τ*). On the other hand, when Δ*τ* ≳ Δ*τ* ^∗^ holds, the exponential factor is diminished and the temporally averaged population of the specialists is suppressed by the term Δ*τ* ^−1^. The discontinuity of the solutions in the two regimes of Δ*τ* regimes in Eq (12) is due to the different approximations of the estimates. We will address this point later.

Consequently, the population size at nutrient supply events (*X*_*i*_(*τ*_*k*_); for1 *k* ≥ 1, *i* = 1, 2 and 3) correspondingly declined. Because fewer cells compete for incoming nutrients, the relative nutrient supply *per capita* increases, and consequently, the duration of nutrient availability after the nutrient supply event (*T* ^+^) is prolonged. As the population size increases exponentially as a function of *T* ^+^, the specialist, which has the highest growth rate among the three phenotypes, exhibits the most pronounced increase. The exponential dependence of the specialists on Δ*τ* is expressed by the factor exp(*c*_2_Δ*τ*).

Once Δ*τ* becomes sufficiently large (Δ*τ* ≳ Δ*τ* ^∗^), total population size decreases. In this regime, the difference between the generalist and specialist populations at nutrient supply events is smaller than that in the small-Δ*τ* region. Because of this small difference, a specialist with a higher growth rate than the others gains an early advantage and leaves little opportunity for the other phenotypes to take up nutrients. Note that this fast-growing specialist also declines rapidly owing to its high death rate, and thus, the overall population size is reduced. This baseline level decrease offsets the exponential term, exp(*c*_2_Δ*τ*), in Eq (12). As a result, the mean abundance of specialists no longer rises but instead declines in proportion to Δ*τ* ^−1^.

We also found that ⟨*X*_*s*_⟩ was proportional to the phenotype switching probability *p*. This implies that inflow from the generalist population sustains a baseline number of the specialists; consequently, the specialists exhibit a pronounced increase at nutrient supply events.

The above conclusions pertain solely to the temporal average of population sizes. The crossover between the temporally averaged populations of generalists and specialists observed in Fig 4B is consistent with the results in the previous section, where the dominance of generalists or specialists is determined by the ratio of growth to death rates, *r*_*g*_*/r*_*s*_.

### Variance of nutrient supply interval

Fig 5 shows the temporally averaged population of the generalist and two specialists when Δ*τ* follows a gamma distribution. The representative trajectories for these regimes are shown in Fig 2. In the generalist-dominant condition, the temporally averaged population of specialists tended to be higher than the constant Δ*τ* . An increase was also observed in the generalist population in the specialist-dominant condition; however, it was smaller than that observed in the specialist population. The differing magnitudes of the increase reflect the tendency of each strategy to expand under non-dominant conditions. The number of generalists rarely increased in the specialist-dominant condition (Fig 2B). In contrast, an increase in the specialist population was often observed in the generalist-dominant condition (indicated by the arrows in Fig 2C).

**Fig 5.**
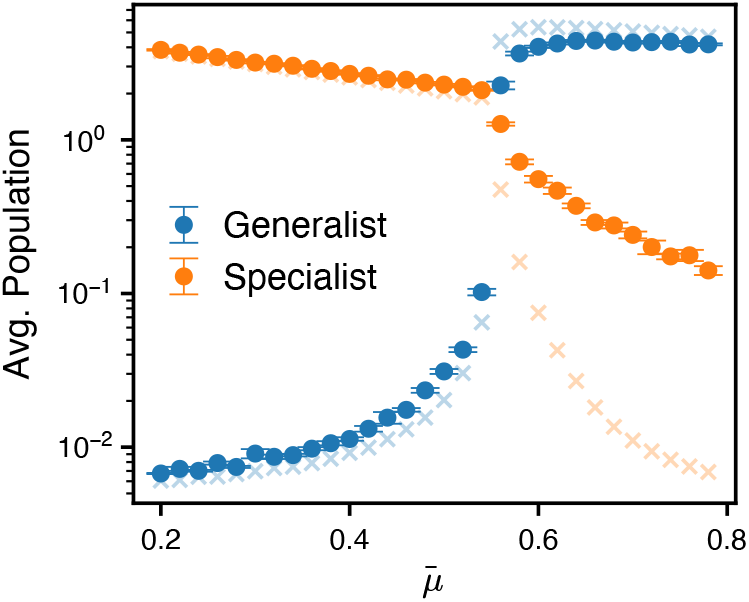
Temporal average of the population with random Δ*τ*. Changes in the temporally averaged populations as the parameter 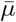 is varied in simulation with one generalist and two specialists. The waiting time for the nutrient supply{ Δ*τ*_*k*_} _*k*≥1_ follows from a gamma distribution, Δ*τ*_*k*_ ∽ Γ(2, 50). We simulated up to *t* = 3.0 10^5^ and averaged the populations after *t* = 2.5 10^5^. For each 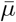, we ran 10 simulations and averaged the results. The error bars indicate the standard error. We also display the results of Δ*τ* = 100 (Fig 3A) in the background (cross marks).

We also performed simulations in which Δ*τ* followed uniform, normal, and gamma distributions with the same expected value 𝔼 [Δ*τ*] = 100, systematically varying the variance of each distribution (S4 Fig). In all cases, we observed that the temporally averaged population of the specialists increased exponentially with the variance in Δ*τ* under the generalist-dominant condition (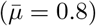), whereas the generalist population increased only slightly under the specialist-dominant condition (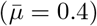).

The analytical estimate also provides insights into the effect of variance in Δ*τ* on population dynamics. When Δ*τ* is a random variable, the temporally averaged population of the generalist and the specialists can be evaluated as the Δ*τ* -weighted expected values of Eq (11) and Eq (12), respectively. Accordingly, the order of ⟨*X*_*s*_⟩ with respect to the increase of Δ*τ* can be estimated as follows:

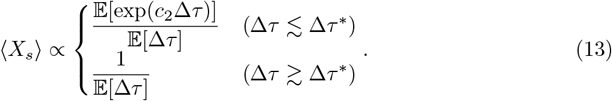

Using the relationship between the moment-generating function and the cumulant-generating function of the random variable Δ*τ*, we can derive the following equation:

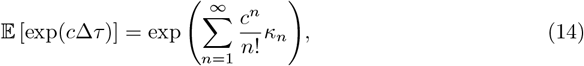

where *κ*_*n*_ is the *n*-th cumulant of Δ*τ, κ*_1_ = 𝔼 [Δ*τ*], *κ*_2_ = Var[Δ*τ*], . Therefore, the temporally averaged population of the specialists increases exponentially with the variance in Δ*τ* when Δ*τ* ≳Δ*τ* ^∗^.

These results imply that in environments where nutrient supply varies, the specialist strategy can temporarily outperform generalists even though it is at a disadvantage in terms of the growth-death ratio. When Δ*τ* follows a probability distribution, occasionally large values are expected. Such infrequent, extended periods without nutrients cause a significant decline in the overall population because cells face prolonged famine. However, immediately following these events, specialists can utilize the next nutrient supply event more rapidly than other phenotypes because of their higher growth rates.

### Diversification of nutrient utilization strategies in multiple environmental conditions

In this section, we increased the number of environmental conditions (*E* = 3 and 4) and observed their dynamics. In natural habitats, there are usually more than two available nutrients, and cells face a resource-use trade-off among multiple nutrients. Under these multi-nutrient conditions, not only two distinct strategies—generalists and specialists—but also a wider range of intermediate phenotypes can be observed. To capture the wider spectrum of possible nutrient utilization strategies, we introduced a finer phenotype space, where growth rates varied smoothly in the constraint defined by Eq (4).

First, to observe how a finer phenotype resolution alters outcomes compared with the discrete two-environment simulations in the previous section, we conducted simulations for two environmental conditions with an increased number of phenotypes (*N* = 51), refining the bins of the phenotype space. As in the simulations with one generalist and two specialists, we set the growth rates of phenotypes in ascending and descending order, and we assumed symmetry between environments A and B. We set the growth rate of phenotype *i* as follows, which is consistent with the resource-use tradeoff.

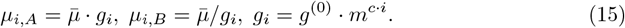

Δ*τ* was set to a constant value, and the environment was changed alternately. Initially, only a generalist-like phenotype was present, with equal growth rates in both environments (*µ*_*i,A*_ = *µ*_*i,B*_).

We simulated the model dynamics for different values of 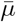, and plotted the temporal average of the population, as shown in Fig 6. Consequently, a continuous shift from a generalist-like strategy to specialist-like strategies was observed by decreasing 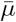. When 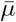 was larger than the branching point at 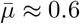 (gray dashed line in Fig 6), the generalist-like strategy becomes dominant. When the value of 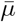 falls below the branching point, the population abundance distribution in the phenotype space becomes bimodal. The phenotypes at the local maxima of the distribution, *α* and *β*, were symmetric between environments A and B (*µ*_*α,A*_ = *µ*_*β,B*_ and *µ*_*α,B*_ = *µ*_*β,A*_). Near the branching point, the two dominant phenotypes showed little difference in growth rate between the two environments. As 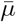 decreases, a specialist-like strategy adapted to only one environment becomes dominant. We also observed a shift from the generalist-like strategy to the two symmetric specialist-like strategies when decreased *b*. As described in the first part of the Results section, we observed a shift from a generalist to specialist dominance. Introducing a continuous phenotype space reveals that this transition does not occur as a sharp transition, but rather unfolds gradually.

**Fig 6.**
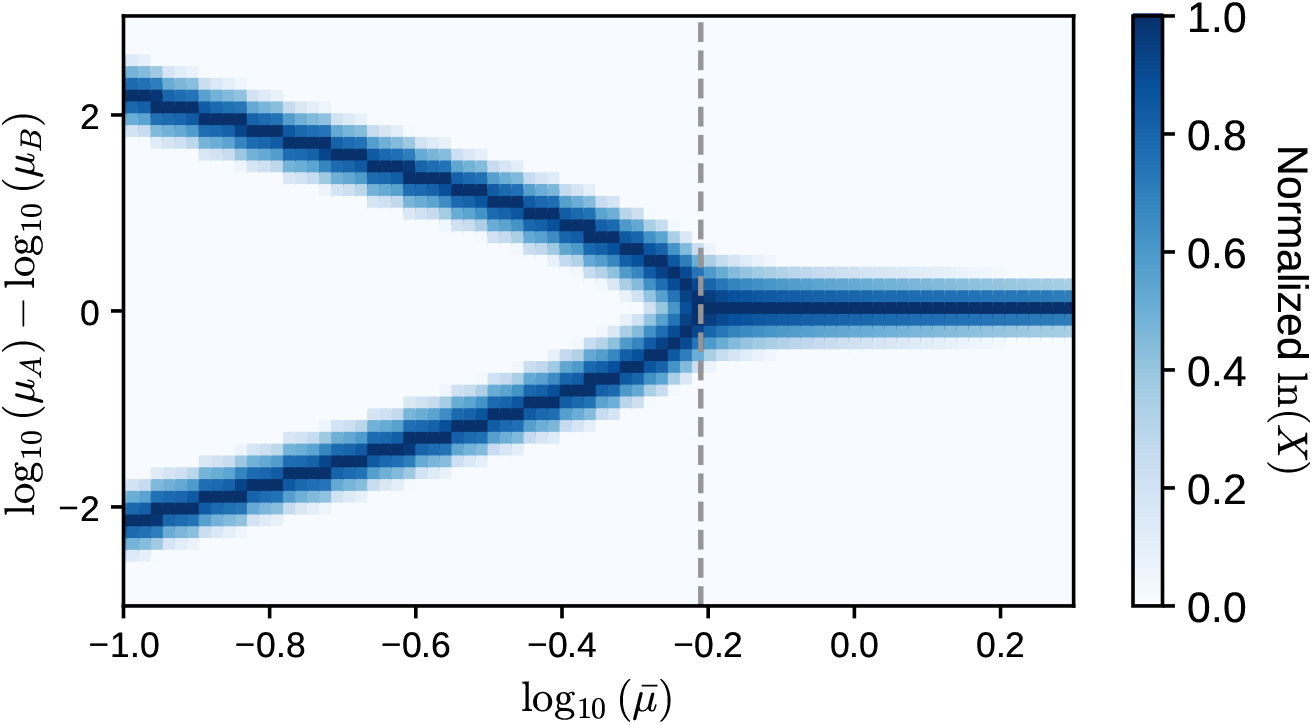
Transition of specialist and generalist. We varied 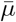 from 10^−1.0^ to 10^0.3^. and ran each simulation until *t* = 2.0 10^5^. Population sizes were time-averaged over the last 20 nutrient supply intervals. For the visibility’s sake, the temporally averaged populations in the logarithmic scale were min-max normalized to [0, 1] in each bin of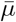. At the 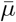 indicated by the gray dashed line, the system transitions from a single dominant phenotype to two co-dominant phenotypes. The parameters are set to *a* = 0.01, *b* = 1, *S*_0_ = 10 and Δ*τ* = 10. The growth rate of each phenotype are set by Eq (15), where *m* = 2, *c* = 0.2, *g*^(0)^ = *m*^*c*(*N*+1)*/*2^. varying in increments of 0.2.

Next, we conducted simulations under three environmental conditions (*E* = 3) to observe the dynamics of the system under more than two environmental conditions. As in the setup above, we discretized the phenotype space 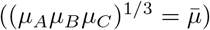 using finely spaced bins. In the initial state, only a single phenotype exists, which satisfies the condition that all the growth rates in different environments are equal.

In the simulations, the dominant strategy changed at a certain value of 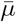, as in the simulation with the two environments. Fig 7 shows the temporal average of the population of each phenotype in one cycle after the system was developed for a sufficiently long period to reach the steady population level on average. As shown in Fig 7A and B, the three peaks stably coexisted. Each peak is located at the position where the growth rates in two of the three environments are the same and higher (e.g., *µ*_*A*_ = *µ*_*B*_ *> µ*_*C*_). Thus, these three phenotypes have specialist-like strategies that are specific to the two environments. Phenotypes that specialize in only one environmental condition were not dominant.

**Fig 7.**
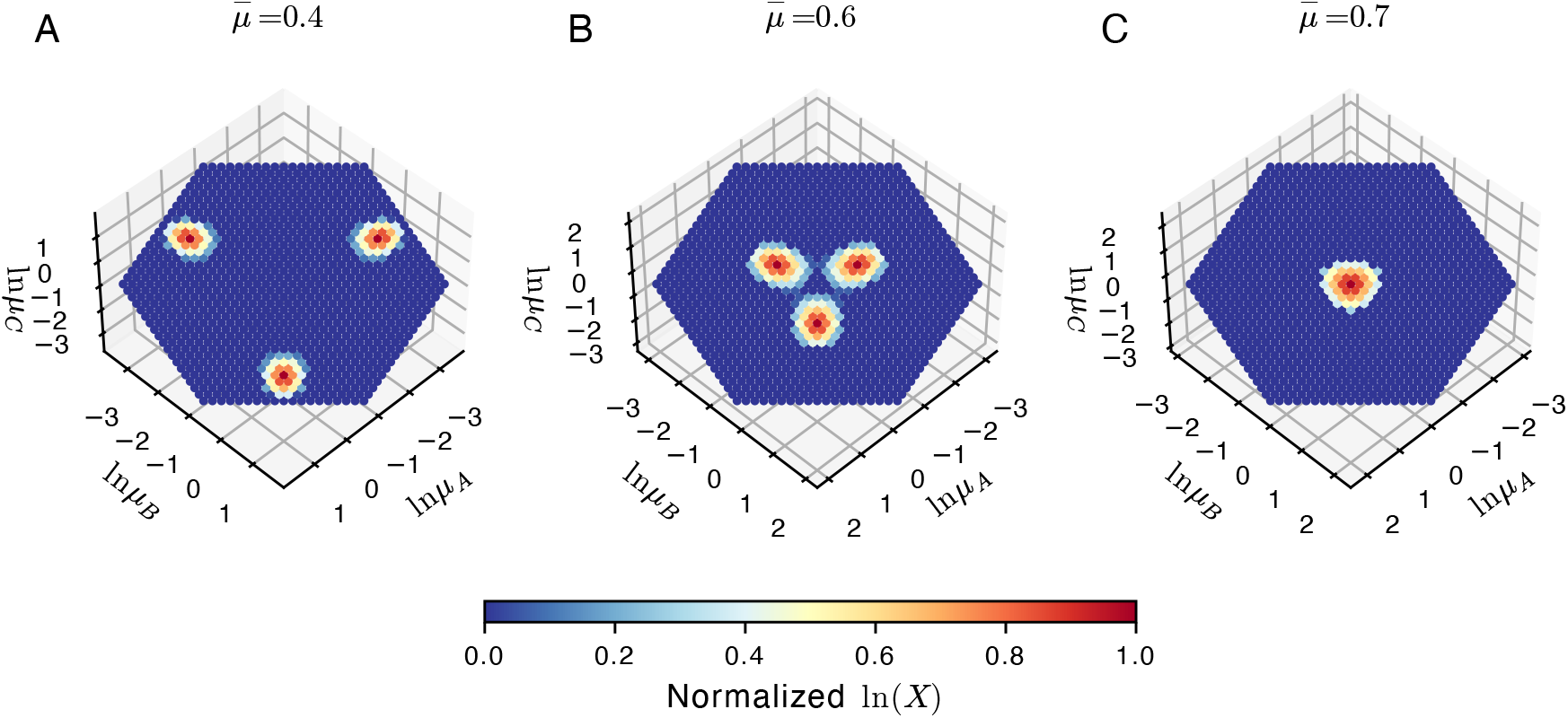
**Examples of the simulation with** *E* = 3. (A)The case with 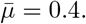 (B) 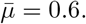. (C) 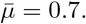. Each dot represents one phenotype. We simulated the population dynamics up to *t* = 2.0 *×* 10^5^. Population sizes were time-averaged over the last 30 nutrient supply intervals. For each simulation, the logarithm of the temporally averaged population in each bin, normalized from 0 to 1, is plotted. Phenotypes with higher populations are marked in red. The parameters are set to *a* = 0.01, *b* = 1, *S*_0_ = 10 and Δ*τ* = 10. The growth rate of each phenotypes in each environment was set as 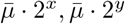 and 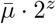, where *x* + *y* + *z* = 0 holds. Phenotypes can switch only between adjacent bins, with a transition probability of 10^−4^.

Simulations were also conducted under four environmental conditions. We observed that when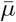 was larger than a certain value, the phenotypes adapted to all four environments were dominant, whereas when 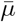 was smaller than a certain value, phenotypes adapted to the three environments emerged. Phenotypes that specialize in one or two environmental conditions were not dominant, as far as we have confirmed. Although we did not conduct experiments with *E* ≥ 5, it is expected that only phenotypes adapted to *E* and *E* − 1 environments will emerge, similar to the case with *E* = 3 and 4.

## Discussion

In the present study, we developed a microbial population dynamics model under the feast-famine cycle to reveal a possible scenario underlying the diverse nutrient utilization strategies of microorganisms in temporally varying environments. In this model, each phenotype is defined by distinct growth rates on different nutrient types, and these rates are constrained by a resource-use trade-off that represents the underlying metabolic functional limitations. Nutrients are supplied to the ecosystem as discrete, stochastic events. Once supplied, they are rapidly consumed by microbes, leading to a nutrient-depleted famine period. The alternation of nutrient-rich and nutrient-depleted conditions leads to a feast-famine cycle. Under the feast-famine cycle, cells face a trade-off between the growth rate in feast and death rate in famine.

In the simplest model, with only three phenotypes and two types of nutrients, we demonstrated that the ratio between the growth rate in feast and death rate in famine determines whether the generalist or specialist phenotype is dominant. When the ratio of the average growth rate to the average death rate exceeds that of its competitors, the phenotype dominates; when it falls below the competitors’ ratio, it is outcompeted. In environments with fluctuating nutrient levels, cells require the regulation of both growth under nutrient-rich conditions and survival under nutrient-poor conditions. However, regulation is not performed instantaneously when the nutrient level changes. For instance, the stress response system should be activated prior to nutrient depletion to achieve high survivability during the famine period at the expense of the growth rate. In our model, this trade-off between growth and survival was effectively incorporated into the growth-death trade-off.

To determine if the above trade-offs lead to the emergence of generalist and specialist strategies, we examined the model dynamics by varying two parameters: 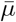 and *b*. We observed that both strategies can be dominant depending on the strength of the growth-death trade-off, even though the metabolic utilization trade-off between different nutrients is fixed to a convex function. While previous theoretical studies have suggested that strong trade-offs tend to favor specialists and limit the emergence of generalists, our results indicate that the outcome can depend on other physiological constraints [16, 38, 39]. However, a situation in which only the functional form of the trade-off type sets the emerging strategy would be contradictory. Indeed, empirical studies have shown that changing only the frequency of environmental fluctuations can reverse which strategy has a higher fitness [24, 27].

The functional form of the resource-use tradeoff is challenging for cells to change. Resource-use trade-offs may be largely influenced by the metabolic reaction systems of the cells. Such trade-offs derived from metabolism are expected to be rigid, as metabolic functions operate under biochemical and thermodynamic constraints, making metabolic innovations, such as the acquisition of new metabolic pathways challenging [46]. Central carbon metabolism, such as glycolysis and the TCA cycle, is evolutionarily conserved across a wide range of organisms [47, 48]. These metabolic pathways constrain key processes such as glycolysis/gluconeogenesis and carbon source co-utilization, and thus represent a key factor shaping resource-use trade-offs [11, 49, 50].

In contrast, the growth-death trade-off may exhibit greater plasticity than the resource-use trade-off. For instance, *rpoS* mutations are commonly found in both laboratory and natural populations [33]. This suggests that the strength of the growth-death tradeoff may evolve through single mutations in regulatory genes.

Taken together, our results suggest that flexible regulation of stress tolerance can compensate for rigid metabolic constraints, allowing microbial communities to toggle between generalist and specialist strategies in response to temporally varying conditions, even when the underlying resource-use trade-off remains unchanged.

We also investigated the effect of the nutrient-supply interval Δ*τ* on population dynamics. We found that the temporal average of specialists increased as the mean and the variance of Δ*τ* increased under the generalist-dominant conditions. This increase can be attributed to the prolonged nutrient-supply intervals (large Δ*τ*) that transiently depress the total population, thereby granting the faster-growing specialist strategy a temporary competitive advantage.

We also found that the temporally averaged population of the specialists was proportional to the phenotype switching probability *p*. This is because the specialist population was sustained by the continuous inflow of mutants originating from the generalist lineage. Because the generalist phenotype—–endowed with a high

growth-death ratio—–constitutes an evolutionarily stable strategy, it persistently acts as a “source,” supplying rare specialist lineages that are unstable yet periodically advantageous (or “sink”). Such source-sink dynamics involving generalist and specialist strategies have been discussed in both theoretical and empirical studies [51–54]. These studies suggest that such dynamics can serve as key mechanisms for the long-term maintenance of diversity in ecological communities.

Our study identifies a key factor that promotes the emergence of diverse resource-use strategies under temporally fluctuating conditions, and provides insight into the broader mechanisms that drive ecosystem diversification.

## Supporting information

**Fig S1.**
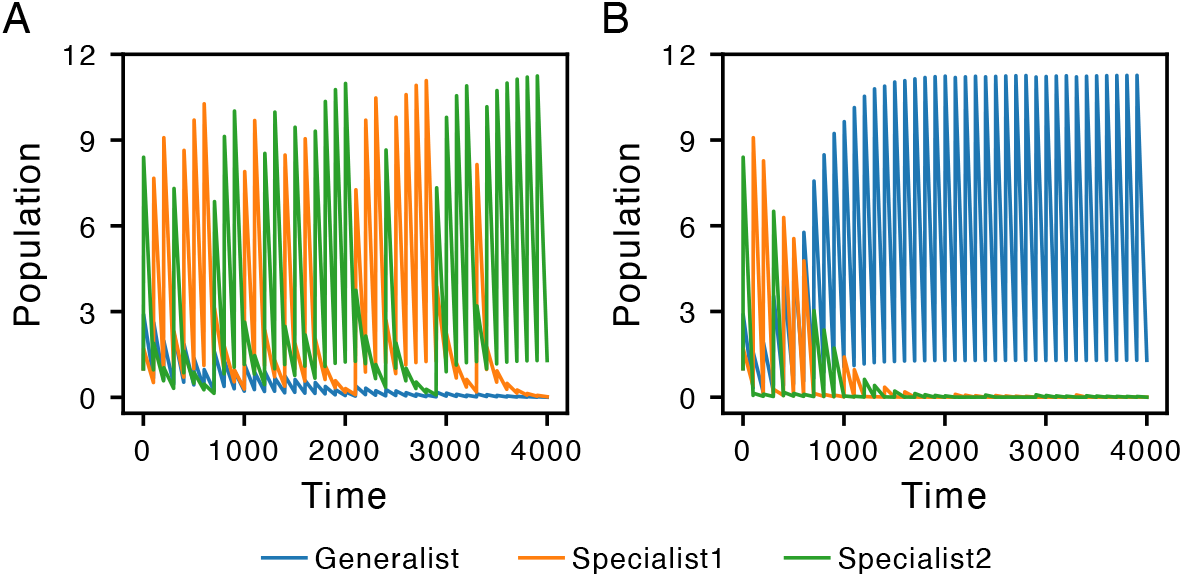
Time series plot of the population with constant Δ*τ*. Examples of the simulation under the condition that nutrient supply events occur at regular intervals, Δ*τ* = 100. (A)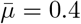, (B) 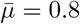. The other parameters were the same as those in Fig 2.

**Fig S2.**
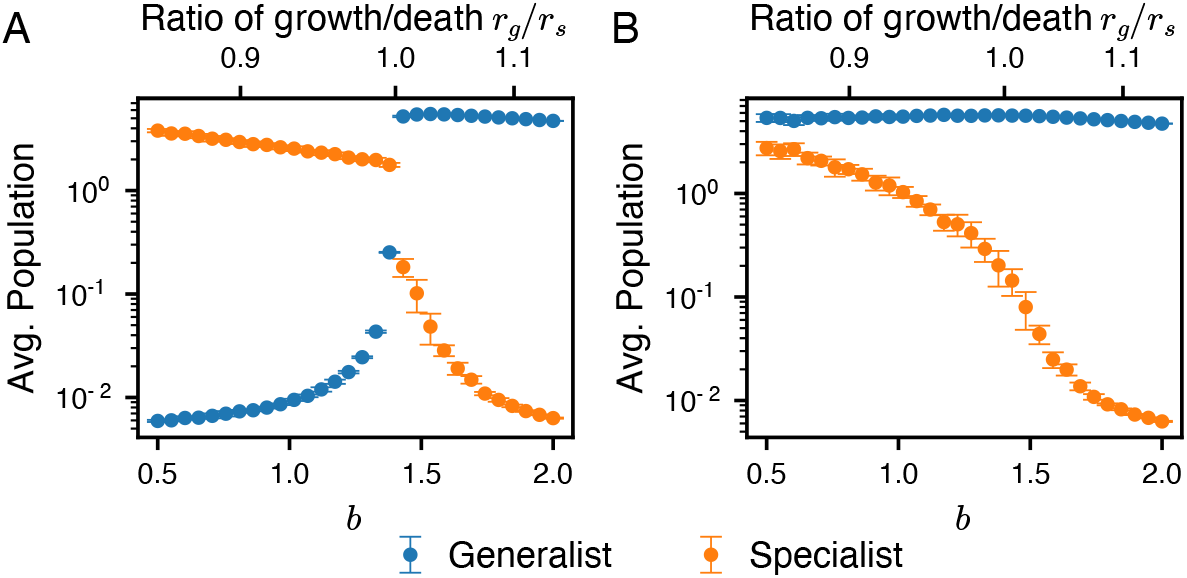
Temporal average of the population. Changes in the temporally averaged populations as parameter *b* was varied in the simulation with (A) *E* = 2, *n* = 3 and (B) *E* = 2, *n* = 2. The geometric average of the growth rate 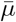 was set to 0.4. The other settings were the same as those in Fig 3.

**Fig S3.**
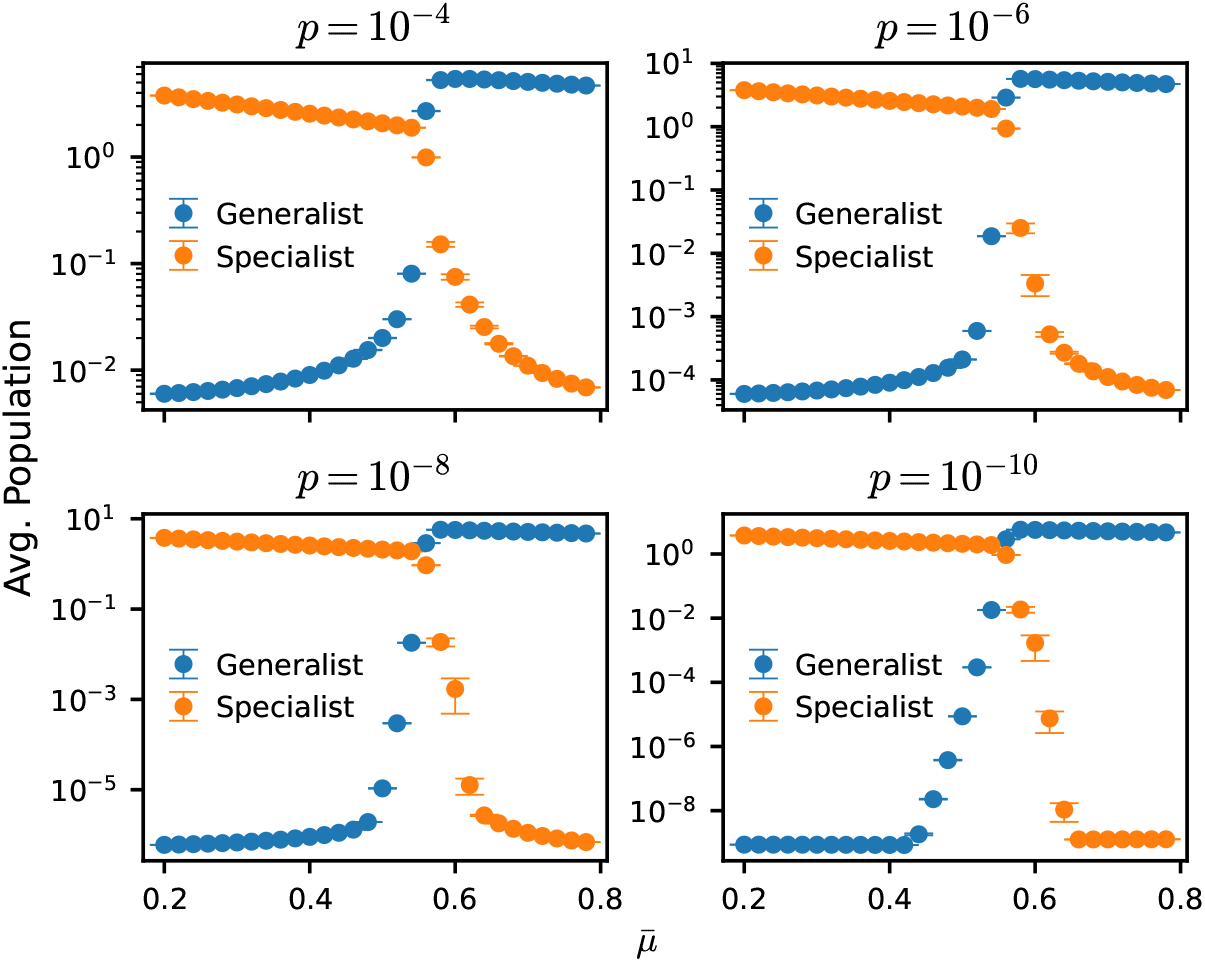
Effect of phenotype switching probability. Temporal average of the population with different phenotype switching probability between generalists and specialists, *p*. Simulations were conducted with *p* = 10^−4^, 10^−6^, 10^−8^ and 10^−10^. The other parameter settings were the same as those in Fig 3.

**Fig S4.**
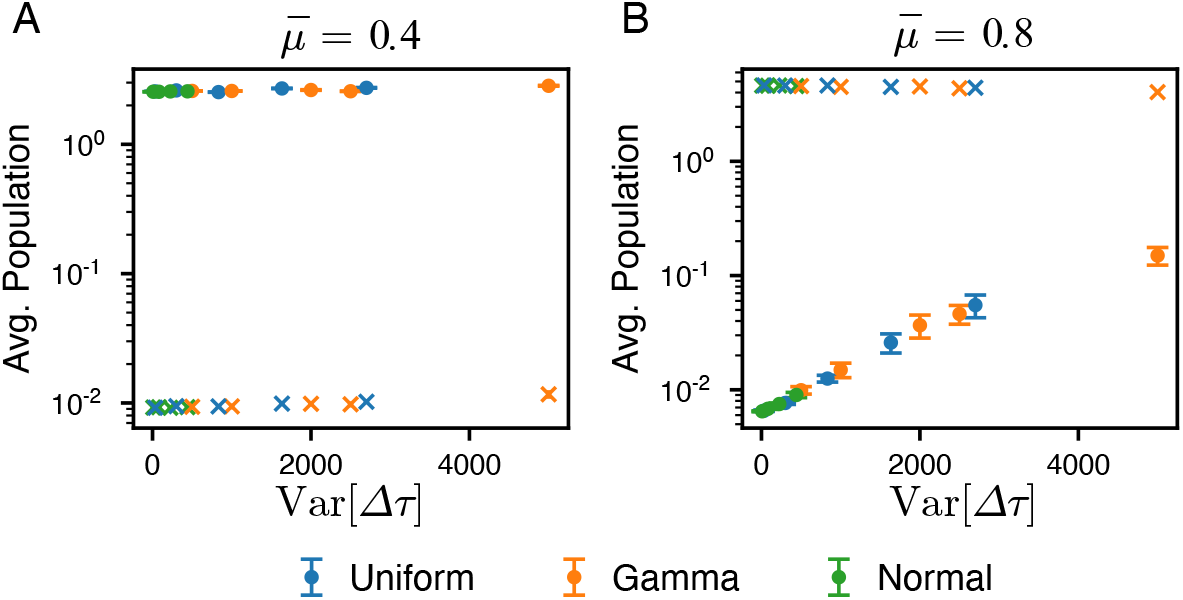
Effect of Δ*τ* variance on the temporal average of the population. Temporal average of the population with different variances in Δ*τ* . We performed simulations with Δ*τ* following uniform, normal and gamma distributions. The dots represent the temporal average of the population of specialists, and the crosses represent those of generalists. The geometric mean of the growth rate 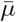differed between the two figures: 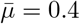 in A and 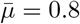 in B. For each of the three distribution families, we generated several variants with the same mean (𝔼 [Δ*τ*] = 100) but different variances and ran the simulations. Uniform distributions were sampled from *U* (100 − *d*, 100 + *d*); for *d* = 10, 30, 50, 70 and 90. Normal distributions were sampled from𝒩 (100, *σ*^2^); for *σ* = 3, 9, 15 and 27. Gamma distributions were sampled from Γ(*k*, 100*/k*); for *k* = 2, 4, 5, 10 and 20. For each distribution, we performed 20 simulations and averaged the results. The error bars indicate standard deviation. All the other parameters were identical to those used in Fig 2.

### S1 Text

**Details of analytical calculations**.

## Acknowledgements

This work was supported by JSPS KAKENHI (Grant Numbers JP22H05403 and JP25H01390 to Y.H.; 22K21344 and 23K27164 to C. F.).

## Notes

### Competing Interest Statement

The authors have declared no competing interest.

### Summary of Updates

Results expanded and clarified; supplementary information added

