## Supplementary Text for "Population dynamics of generalist and specialist strategies under feast–famine cycles"

#### Contents


1 The specific choice of resource-use trade-off function did not alter the result 1

2 Evolutionary invasion analysis. 2

3 Analytical solution for the temporally averaged population under generalist-dominant condition 4

4 Order estimation of the temporally averaged population 8


#### 1 The specific choice of resource-use trade-off function did not alter the result

In the main body of the paper, we imposed the following constraint on the resource-use trade-off:

$$\left(\prod_{e=1}^{E} \mu_{i,e}\right)^{\frac{1}{E}} = \bar{\mu}, \tag{1}$$

where $\mu_{i,e}$ denotes the growth rate of phenotype $i$ under environment $e$, and $\bar{\mu}$ is the prescribed geometric mean growth rate across all $E$ resources.

To test the robustness of our findings, we further examined a canonical nonlinear convex trade-off function originally introduced in a previous study [1]:

$$\sum_{e=1}^{E} \mu_{i,e}^{\gamma} = m, \tag{2}$$

where $\gamma$ and $m$ are the positive constants. Parameter $\gamma$ modulates the curvature of the trade-off function, governing whether it is convex ($\gamma < 1$) or concave ($\gamma > 1$), while $\gamma = 1$ yields a linear trade-off.

We adopted $\gamma = 0.8$ in the following simulations. As described in the main text, we performed simulations with three phenotypes (phenotype 1,2, and 3) and two environmental conditions (environment A and B). The growth rates of the three phenotypes were set as follows:

$$\mu_{1,A} = \mu_{3,B} = (0.9m)^{\frac{1}{\gamma}}, \mu_{2,A} = \mu_{2,B} = (0.5m)^{\frac{1}{\gamma}}, \mu_{3,A} = \mu_{1,B} = (0.1m)^{\frac{1}{\gamma}}. \tag{3}$$

Phenotype 2 has a generalist-like strategy, while phenotype 1 and 3 are specialists adapted to Environment A and B, respectively.

Using this setup, we ran simulations and obtained results that were consistent with the results of the resource-use trade-off specified by Eq (1) (see main text). Namely,

- The dominant phenotype was determined by the growth-death ratio (Fig A).
- Increasing the mean and variance of $\Delta\tau$ increases the time average of the population of specialists (Fig B and Fig C).

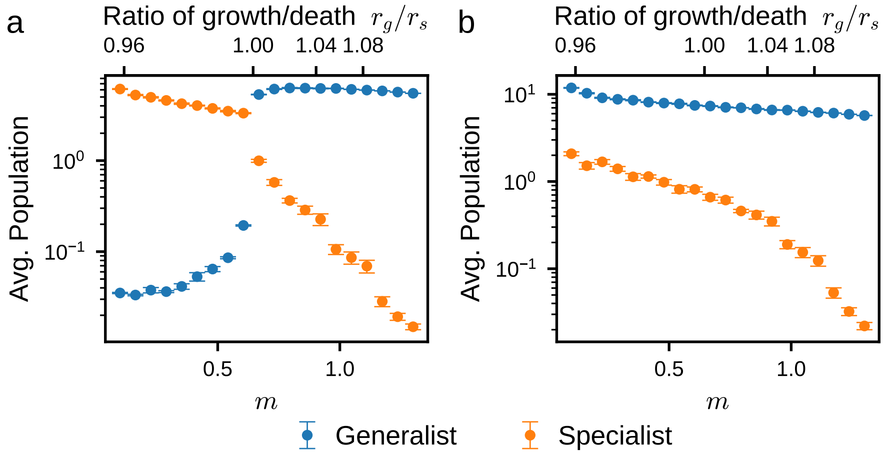


**Fig A. Temporal average of the population with constant $\Delta\tau$.** The average population numbers are shown as a function of the parameter $m$. The model with one generalist and two specialists is simulated for (a), while there is only a single specialist adapted to environment A for (b). We simulated up to $t = 3.0 \times 10^5$ and averaged the populations after $t = 2.5 \times 10^5$. For each $m$, we ran 10 simulations and calculated the average population. The error bar indicates the standard error. All other parameters were identical to those used in Fig 2 in the main text.

#### 2 Evolutionary invasion analysis.

To clarify why the transition occurred at $r_g/r_s = 1$, we estimated this transition point analytically. We considered a situation in which only one phenotype exists in the system and is in a steady state and examined whether the invasive phenotype can be fixed. We also consider that nutrient supply events occur at regular intervals; and thus, $\Delta\tau$ is constant.

First, let $X_i^{(k)}$ be the population of phenotype $i$ at the $k$-th nutrient supply event ($X_i^{(k)} = X_i(\tau_k)$). The change in population size caused by a nutrient supply event is determined by an increase in population in the feast phase and a decrease in population in the famine phase. Thus, $X_i^{(k+1)}$ can be calculated as

$$X_i^{(k+1)} = X_i^{(k)} \exp\left(\mu_{i,e_k} T_k^+ - \gamma_{i,e_k}(\Delta\tau - T_k^+)\right). \tag{4}$$

$T_k^+$ denotes duration of nutrient availability after $k$-th nutrient supply event. We define $f_{i,k}$ as the logarithm of the ratio of $X_i^{(k+1)}$ to $X_{i+1}^{(k)}$.

$$f_{i,k} = \mu_{i,e_k} T_k^+ - \gamma_{i,e_k}(\Delta\tau - T_k^+). \tag{5}$$

When $f_{i,k}$ is positive, the population of phenotype $i$ increases, whereas when $f_{i,k}$ is negative, the population decreases. Because the two environments come at random with equal probability, the average value of fitness $f_i$ is

$$f_i = \langle f_{i,k} \rangle = \mu_i T^+ - \gamma_i(\Delta\tau - T^+), \tag{6}$$

where $\mu_i = \dfrac{\mu_{i,A} + \mu_{i,B}}{2}, \gamma_i = \dfrac{\gamma_{i,A} + \gamma_{i,B}}{2}$. Suppose that only one phenotype (phenotype $\alpha$) exists, and the system is in a steady state; ($X_\alpha^k = X_\alpha^{k+1} = X_\alpha^{st}$), $f_\alpha = 0$ holds.

$$f_\alpha = \mu_\alpha T^+ - \gamma_\alpha(\Delta\tau - T^+) = 0. \tag{7}$$

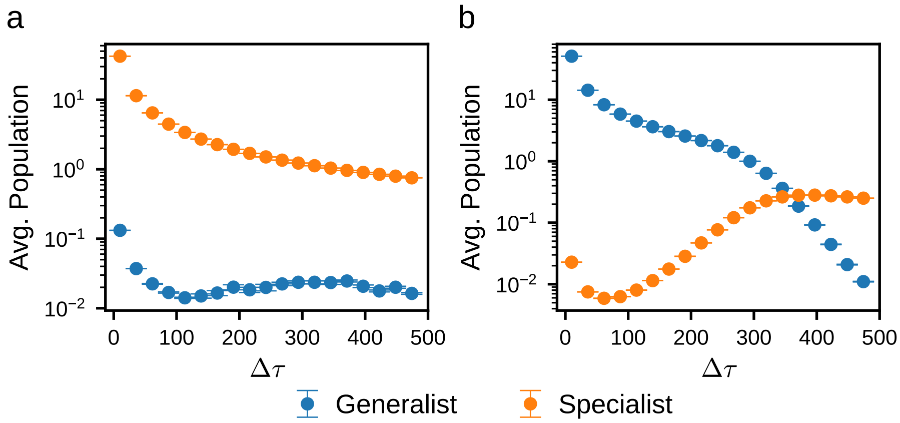


**Fig B. Temporal average of the population as a function of $\Delta\tau$.** Changes in the average populations as the parameter $\Delta\tau$ is varied in simulation with one generalist and two specialists. The case with $m = 0.5$ is shown in (a) and the case with $m = 1.5$ is shown in (b). We simulated up to $t = 3000 \times \Delta\tau$ and averaged the populations after $t = 2500 \times \Delta\tau$. $\Delta\tau$ is constant value. For each $\Delta\tau$, we ran 10 simulations and calculated the average population. The error bars indicate the standard error. All other parameters were identical to those used in Fig 4 in the main text.

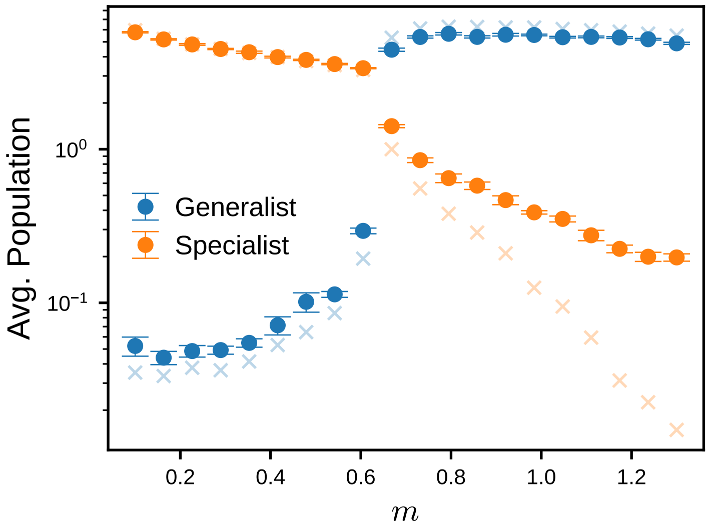


**Fig C. Temporal average of the population with random $\Delta\tau$.** Changes in the average populations as the parameter $m$ is varied in simulation with one generalist and two specialists. The waiting time for the nutrient supply $\{\Delta\tau_k\}_{k\geq 1}$ are sampled from a gamma distribution, $\Delta\tau_k \sim \Gamma(2, 50)$. We simulated up to $t = 3.0 \times 10^5$ and averaged the populations after $t = 2.5 \times 10^5$. For each $m$, we ran 10 simulations and calculated the average population. The error bars indicate the standard error. We also display the results of $\Delta\tau = 100$ (Fig A(a)) in the background (cross marks).

Then, we consider that a small number of another phenotype (phenotype $\beta$) invades the steady state of phenotype $\alpha$. The number of phenotype $\beta$ is sufficiently small such that the change in the value of $T^+$ is negligible. Thus, $f_\beta$ is given by

$$f_\beta \quad = \mu_\beta T^+ - \gamma_\beta(\Delta\tau - T^+). \tag{8}$$

By Solving equation Eq (7) for $T^+$, we get

$$T^+ = \frac{\gamma_\alpha}{\mu_\alpha + \gamma_\alpha}\Delta\tau. \tag{9}$$

Inserting Eq (9) into Eq (8), we get

$$f_\beta = \frac{\Delta\tau}{\mu_\alpha + \gamma_\alpha}\left(\mu_\beta\gamma_\alpha - \mu_\alpha\gamma_\beta\right). \tag{10}$$

Therefore, when $\frac{\mu_\beta}{\gamma_\beta} > \frac{\mu_\alpha}{\gamma_\alpha}$, $f_\beta$ becomes positive so that phenotype $\beta$ can be fixed. On the other hand, when $\frac{\mu_\beta}{\gamma_\beta} < \frac{\mu_\alpha}{\gamma_\alpha}$, $f_\beta$ becomes negative and phenotype $\beta$ is excluded and the steady state with only phenotype $\alpha$ is stable.

### 3 Analytical solution for the temporally averaged population under generalist-dominant condition

To explain the increase in specialists under generalist-dominant conditions, we analytically estimated the temporally averaged population using a simplified model under three assumptions.

1. Specialist populations are small enough that phenotypic change from specialist to generalist is negligible.
2. Two environments A and B change alternately and $\Delta\tau$ is constant.
3. The dynamics settle into a periodic solution. The generalist population show no net change between the nutrient supply events. Specialist populations show no net change between the two consecutive supply events.

There are three phenotypes (phenotype 1, 2 and 3) and two environmental conditions (environment A and B). Phenotype 1 is a specialist in environment A, phenotype 3 is a specialist in environment B, and phenotype 2 is a generalist. We assume that the two environments A and B are symmetric. Thus, we set the growth rates as follows using the three variables $\mu_s^+$, $\mu_s^-$, and $\mu_g$ ($\mu_s^- < \mu_g < \mu_s^+$),

$$\mu_{1,A} = \mu_{3,B} = \mu_s^+,\ \mu_{1,B} = \mu_{3,A} = \mu_s^-,\ \mu_{2,A} = \mu_{2,B} = \mu_g. \tag{11}$$

Similarly, we set the death rates as

$$\gamma_{1,A} = \gamma_{3,B} = \gamma_s^+,\ \gamma_{1,B} = \gamma_{3,A} = \gamma_s^-,\ \gamma_{2,A} = \gamma_{2,B} = \gamma_g, \tag{12}$$

where $\gamma_s^- < \gamma_g < \gamma_s^+$ holds true. Phenotype 2 grows at a rate $\mu_g$ and switches to phenotype 1 and 3 at a rate $p$, respectively. Therefore, generalist population $X_2$ is as follows:

$$\frac{dX_2}{dt} = \begin{cases} (1-2p)\mu_g X_2 & (\tau_k < t < \tau_k + T_k^+) \\ -\gamma_g X_2 & (\tau_k + T_k^+ < t < \tau_{k+1}) \end{cases}, \tag{13}$$

where $T_k^+$ is the duration of nutrient availability after the $k$-th nutrient supply event. The specialist population increases through the sum of its intrinsic growth and inflow of individuals from the generalist population. When the $k$-th nutrient supply event places the system in environment $e$, specialist populations $X_1$ and $X_3$ follow the following equations:

$$\frac{dX_i}{dt} = \begin{cases} \mu_{i,e}X_i + p\mu_g X_2 & (\tau_k < t < \tau_k + T_k^+) \\ -\gamma_{i,e}X_i & (\tau_k + T_k^+ < t < \tau_{k+1}) \end{cases}. \tag{14}$$

Because each supply event resets the nutrient concentration to a fixed level $S_0$, which is subsequently depleted by the three phenotypes, the nutrient dynamics are governed by the following equation.

$$S(\tau_k) = S_0,\ \frac{dS}{dt} = -\left(\mu_g X_2 + \mu_{1,e}X_1 + \mu_{3,e}X_3\right) \quad (\tau_k < t < \tau_k + T_k^+). \tag{15}$$

First, we calculated the temporally averaged population of the generalist. By solving Eq (13), we obtain:

$$X_2(\tau_k + t') = \begin{cases} X_2(\tau_k)\exp(\mu_g^* t') & (0 < t' < T_k^+) \\ X_2(\tau_k)\exp(\mu_g^* T_k^+ - \gamma_g(t' - T_k^+)) & (T_k^+ < t' < \Delta\tau) \end{cases}, \tag{16}$$

where $\mu_g^* = (1-2p)\mu_g$ is the effective growth rate of the generalist.

Assuming that there is no change in the population between nutrient supply events, we have

$$X_2(\tau_k) = X_2(\tau_{k+1}) = X_g^*.$$

Accordingly, the condition

$$\exp(\mu_g^* T_k^+ - \gamma_g(\Delta\tau - T_k^+)) = 1 \tag{17}$$

holds. Thus, $T_k^+$ is given by

$$T_k^+ = \frac{\gamma_g}{\mu_g^* + \gamma_g}\Delta\tau. \tag{18}$$

By integrating $X_2$ from $\tau_k$ to $\tau_{k+1}$, we can calculate the temporally averaged population of the generalist from $\tau_k$ to $\tau_{k+1}$, $\langle X_2\rangle_k$:

$$\begin{aligned} \langle X_2\rangle_k &= \frac{1}{\Delta\tau}\left(\int_{\tau_k}^{\tau_k+T_k^+} X_2(t)\,dt + \int_{\tau_k+T_k^+}^{\tau_{k+1}} X_2(t)\,dt\right) \\ &= X_g^*\left(\frac{1}{\mu_g^*} + \frac{1}{\gamma_g}\right)\left(\exp(\mu_g^* T_k^+) - 1\right), \end{aligned} \tag{19}$$

where we utilized Eq (17) to obtain the final equality.

Next, we calculated the population of specialists. Because we assumed that environment A and B are symmetric and change alternately, the periodic solution of the specialist populations $X_1$ and $X_3$ are symmetric with respect to the two environmental conditions. That is, $X_1(t' + \tau_k) = X_3(t' + \tau_{k+1})$ and $X_3(t' + \tau_k) = X_1(t' + \tau_{k+1})$ hold for all $t'$ satisfying $0 < t' < \Delta\tau$. Therefore, it is sufficient to solve solely for $X_1$.

To simplify the calculations, we introduced the dimensionless quantity $R = X_1/X_2$ and assume that $p$ is negligible. Because $\dot{R} = \frac{\dot{X}_1}{X_2} - R\frac{\dot{X}_2}{X_2}$, it follows from Eq (13) and Eq (14) that

$$\frac{dR}{dt} = \begin{cases} \left(\mu_{1,e} - \mu_g^*\right)R + p\mu_g & (\tau_k < t < \tau_k + T_k^+) \\ -\left(\gamma_{1,e} - \gamma_g\right)R & (\tau_k + T_k^+ < t < \tau_{k+1}) \end{cases}. \tag{20}$$

By solving Eq (20), we obtain

$$R(\tau_k + t') = \begin{cases} R(\tau_k)\exp(\Delta\mu_e t') + \frac{p\mu_g}{\Delta\mu_e}\left(\exp(\mu_e t') - 1\right) & (0 < t' < T_k^+) \\ R(\tau_k + T_k^+)\exp\left(-\Delta\gamma_e(t' - T_k^+)\right) & (T_k^+ < t' < \Delta\tau) \end{cases}, \tag{21}$$

where $\Delta\mu_e = \mu_{1,e} - \mu_g^*$ and $\Delta\gamma_e = \gamma_{1,e} - \gamma_g$.

Assuming that the dynamics settle into a periodic orbit, the population of phenotype 1 returns to its original value after experiencing environment A and B. Therefore, $R(\tau_k) = R(\tau_{k+2})$ holds. In the following, we now solve for $R(\tau_k)$ satisfying this equation.

We introduced the discrete map between $R(\tau_k)$ and $R(\tau_{k+1})$, $F_e$ where $F_e(R(\tau_k)) = R(\tau_{k+1})$ holds. From Eq (21), $F_e$ can be expressed as a linear function of $R$

$$F_e(R(\tau_k)) = E_e R(\tau_k) + D_e, \tag{22}$$

where $E_e = \exp\left(\Delta\mu_e T_k^+ - \Delta\gamma_e(\Delta\tau - T_k^+)\right)$ and $D_e = \frac{p\mu_g}{\Delta\mu_e}\left(E_e - \exp(-\Delta\gamma_e(\Delta\tau - T_k^+))\right)$.

When the $k$-th environment is A, the $(k+1)$-th environment is B, and $R(\tau_{k+2})$ can be calculated as:

$$R(\tau_{k+2}) = F_B(F_A(R(\tau_k))) = E_A E_B R(\tau_k) + D_A E_B + D_B. \tag{23}$$

By setting $R(\tau_k) = R(\tau_{k+2}) = R_A^*$, we obtain

$$R_A^* = \frac{D_A E_B + D_B}{1 - E_A E_B}. \tag{24}$$

Analogously, when the $k$-th environment is B, we can obtain $R_B^*$ by the same procedure as in Eq (23) and Eq (24).

$$R_B^* = \frac{D_B E_A + D_A}{1 - E_A E_B}. \tag{25}$$

Assuming that the system has settled into a periodic solution, Eq (21) can be reformulated using Eq (24) and Eq (25), as follows:

$$R(\tau_k + t') = \begin{cases} R_e^*\exp(\Delta\mu_e t') + \frac{p\mu_g}{\Delta\mu_e}\left(\exp(\Delta\mu_e t') - 1\right) & (0 < t' < T_k^+) \\ R_e^*(T_k^+)\exp(-\Delta\gamma_e(t' - T_k^+)) & (T_k^+ < t' < \Delta\tau) \end{cases}, \tag{26}$$

where $R_e^*(T_k^+) = R_e^*\exp(\Delta\mu_e t') + \frac{p\mu_g}{\Delta\mu_e}\left(\exp(\Delta\mu_e t') - 1\right)$.

The temporally averaged population of phenotype 1 from $t = \tau_k$ to $\tau_{k+1}$ is obtained by integrating $RX_2$ from $t = \tau_k$ to $\tau_{k+1}$ in each environment $e$, and then averaging the values over the two environments, that is

$$\langle X_1 \rangle_k = \frac{1}{2\Delta\tau}\sum_{e=A,B}\left(\int_{\tau_k}^{\tau_k + T_k^+} R(t; e_k = e) X_g(t)\, dt + \int_{\tau_k + T_k^+}^{\tau_{k+1}} R(t; e_k = e) X_g(t)\, dt\right). \tag{27}$$

The first and the second terms of the integral are given by

$$\begin{aligned} \int_{\tau_k}^{\tau_k + T_k^+} R(t; e_k = e) X_g(t)\, dt = {} & \frac{X_g^* R_e^*}{\mu_{1,e}}\left(\exp(\mu_{1,e} T_k^+) - 1\right) \\ & + \frac{X_g^* p\mu_g}{\Delta\mu_e}\left(\frac{\exp(\mu_{1,e} T_k^+) - 1}{\mu_{1,e}} - \frac{\exp(\mu_g^* T_k^+) - 1}{\mu_g^*}\right), \end{aligned} \tag{28}$$

and

$$\int_{\tau_k+T_k^+}^{\tau_{k+1}} R(t; e_k = e) X_g(t)\, dt = X_g^* \exp(\mu_g^* T_k^+) R_e^*(T_k^+) \frac{1-\exp(-\gamma_{1,e}(\Delta\tau - T_k^+))}{\gamma_{1,e}}, \tag{29}$$

respectively.

By inserting Eq (28) and Eq (29) into Eq (27), we can obtain

$$\langle X_1 \rangle_k = \frac{X_g^*}{2\Delta\tau} \sum_{e=A,B} (\mathcal{R}_e + \mathcal{S}_e), \tag{30}$$

where

$$\mathcal{R}_e = \frac{R_e^*}{\mu_{1,e}} \left(\exp(\mu_{1,e} T_k^+) - 1\right) + \frac{p\mu_g}{\Delta\mu_e} \left( \frac{\exp(\mu_{1,e} T_k^+) - 1}{\mu_{1,e}} - \frac{\exp(\mu_g^* T_k^+) - 1}{\mu_g^*} \right) \tag{31}$$

$$\mathcal{S}_e = \exp(\mu_g^* T_k^+) R_e^*(T_k^+) \frac{1-\exp(-\gamma_{1,e}(\Delta\tau - T_k^+))}{\gamma_{1,e}}. \tag{32}$$

The temporally averaged population of generalists and specialists is given by Eq (19) and Eq (30), respectively: however, constant $X_g^*$ remains undetermined. In the following, we calculate $X_g^*$ using Eq (15).

Because the periodic solutions of the specialist populations are symmetric with respect to environment $A$ and $B$, $\mu_{3,A} X_3 = \mu_{1,B} X_1$ and $\mu_{1,A} X_1 = \mu_{3,B} X_3$ hold true.

Therefore, Eq (15) can be rewritten as

$$\frac{dS}{dt} = -\mu_g X_2 - \mu_{1,A} X_1 - \mu_{1,B} X_1. \tag{33}$$

We integrate Eq (33) from $\tau_k$ to $\tau_k + T_k^+$.

$$\begin{aligned} S_0 &= \mu_g \int_{\tau_k}^{\tau_k+T_k^+} X_2(t)\, dt + \sum_{e=A,B} \mu_{1,e} \int_{\tau_k}^{\tau_k+T_k^+} R(t; e_k = e) X_2(t)\, dt \\ &= \frac{X_g^* \mu_g}{\mu_g^*} \left(\exp(\mu_g^* T_k^+) - 1\right) + \sum_{e=A,B} \mu_{1,e} X_g^* \mathcal{R}_e. \end{aligned} \tag{34}$$

By solving Eq (34) for $X_g^*$, we obtain

$$X_g^* = \frac{S_0}{\frac{\mu_g}{\mu_g^*}\left(\exp(\mu_g^* T_k^+) - 1\right) + \sum_{e=A,B} \mu_{1,e} \mathcal{R}_e}. \tag{35}$$

Combining Eq (19) and Eq (30) and Eq (35), we obtain the temporally averaged population of generalists and specialists.

$$\begin{aligned} \langle X_2 \rangle &= \frac{\sum_k \langle X_2 \rangle_k \Delta\tau}{\sum_k \Delta\tau} \\ &= \frac{1}{\Delta\tau} \frac{S_0 \left(\exp\left(\mu_g^* T_k^+\right) - 1\right)}{\frac{\mu_g}{\mu_g^*}(\exp\left(\mu_g^* T_k^+\right) - 1) + \sum_{e=A,B} \mu_{1,e} \mathcal{R}_e} \left( \frac{1}{\mu_g^*} + \frac{1}{\gamma_g} \right), \end{aligned} \tag{36}$$

$$\begin{aligned} \langle X_1 \rangle &= \frac{\sum_k \langle X_1 \rangle_k \Delta\tau}{\sum_k \Delta\tau} \\ &= \frac{1}{\Delta\tau} \frac{S_0 \sum_{e=A,B} (\mathcal{R}_e + \mathcal{S}_e)}{\frac{\mu_g}{\mu_g^*}(\exp\left(\mu_g^* T_k^+\right) - 1) + \sum_{e=A,B} \mu_{1,e} \mathcal{R}_e}, \end{aligned} \tag{37}$$

where $T_k^+$,$\mathcal{R}_e$ and $\mathcal{S}_e$ are defined by Eq (18), Eq (31) and Eq (32), respectively.

Fig D shows a comparison of the analytical solution with the numerical simulation results. The simulation parameters are identical to those used in Fig 4B in the main text (e.g. $\bar{\mu} = 0.8$). We substituted these parameter values into Eq (36) and Eq (37) to compute the corresponding analytical solutions. The dots indicate the simulation results, and the dashed curves show the analytical solution. Fig D(a) shows simulations in which environments A and B alternate deterministically, while Fig D(b) shows simulations in which the sequence of environments A and B is randomized. In simulations with alternating environments, the analytical solution was in good agreement with the simulation results. In simulations with randomly fluctuating environments, the analytical prediction for the generalist deviates at a large $\Delta\tau$; nonetheless, it still captures the principal qualitative trend.

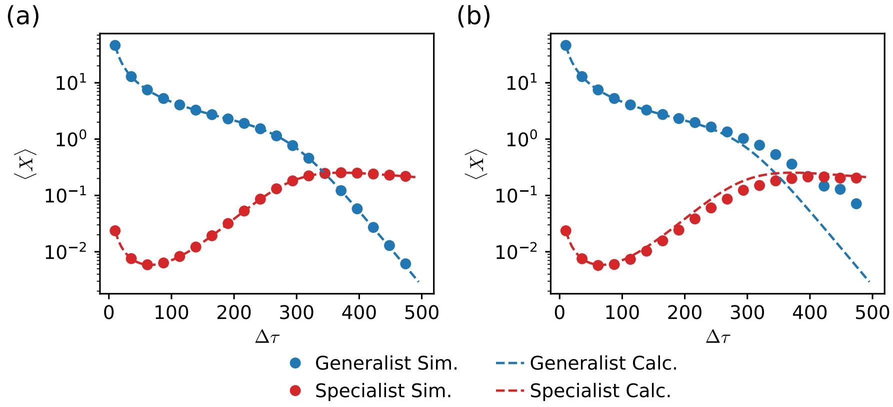


**Fig D. Comparison of the analytical solution with the numerical simulation results.** The dots indicate the simulation results, and the dashed curves show the analytical solution. (a) Simulations in which environments A and B alternate deterministically. (b) Simulations in which the sequence of environments A and B is randomized. The parameters are set to $\mu_{2,A} = \mu_{2,B} = 0.8$, $\mu_{1,A} = \mu_{3,B} = 1.6$, $\mu_{1,B} = \mu_{3,A} = 0.4$, $a = 0.01$, $b = 1$, $p = 10^{-4}$ and $\theta = 10^{-8}$. The dots in (b) are identical to those in Fig 4B in the main text.

### 4 Order estimation of the temporally averaged population

In this section, we estimate the order of the temporally averaged population in response to increasing $\Delta\tau$ using the analytical solution Eq (36) and Eq (37). Based on Eq (18), which shows that $T_k^+ \propto \Delta\tau$, we treated $\mathcal{R}_e$, $\mathcal{S}_e$, and $X_g^*$ as functions of $\Delta\tau$. Fig E(a), (b) and (c) show the analytical solutions $\mathcal{R}_e$, $\mathcal{S}_e$, and $X_g^*$ as functions of $\Delta\tau$, respectively. All three quantities vary exponentially with respect to $\Delta\tau$. We estimated the exponential coefficient and presented them in the figure legend.

$X_g^*$ decreases exponentially with $\Delta\tau$. Two distinct decay regimes are evident: for $\Delta\tau \lesssim 300$ the estimated index is about $-2.6 \times 10^{-2}$, while for $\Delta\tau \gtrsim 300$ it is about $-5.1 \times 10^{-2}$.

When $\mu_g = 0.8$, $\mu_g^* T_k^+ = \frac{\mu_g^* \gamma_g}{\mu_g^* + \gamma_g} \Delta\tau \approx 2.2 \times 10^{-2} \times \Delta\tau$ holds. Hence, for $\Delta\tau \lesssim 300$, the decrease in $X_g^*$ is almost offset by the factor $\exp(\mu_g^* T_k^+)$; consequently, the prefactor $1/\Delta\tau$ becomes the principal driver of the decrease in $\langle X_2 \rangle$. In contrast, for $\Delta\tau \gtrsim 300$,

$\langle X_2 \rangle$ decreases more rapidly owing to attenuation of both the $1/\Delta\tau$ and the exponential function. In summary, $\langle X_2 \rangle$ roughly decreases as follows:

$$\langle X_2 \rangle \propto \begin{cases} \Delta\tau^{-1} & (\Delta\tau \lesssim 300) \\ \Delta\tau^{-1} \exp(-c_1 \Delta\tau) & (\Delta\tau \gtrsim 300) \end{cases} . \tag{38}$$

$c_1$ is a constant value and is approximately $c_1 \approx 2.9 \times 10^{-2}$ (Fig E(d)).

Next, we estimated order of $\langle X_1 \rangle$. For both $\mathcal{R}_e$ and $\mathcal{S}_e$, the exponent is larger when $e = A$ than when $e = B$. Because $\mathcal{S}_A$ is approximately two orders of magnitude larger than $\mathcal{R}_A$, the variation in $\langle X_1 \rangle$ is governed primarily by the terms $X_g^*$, $1/\Delta\tau$ and $\mathcal{S}_A$. For $\Delta\tau \lesssim 300$, the positive contribution from the increase in $\mathcal{S}_A$ outweighs the negative contributions from the decrease in $X_g^*$. However, for $\Delta\tau \gtrsim 300$, both contribution are almost offset, and only the negative contribution from $1/\Delta\tau$ dominates. Furthermore, noting that both $\mathcal{R}_e$ and $\mathcal{S}_e$ are proportional to the phenotype switching probability $p$ ($\mathcal{R}_e, \mathcal{S}_e \propto p$), we can express the temporally averaged population of phenotype 1 as follows:

$$\langle X_1 \rangle \propto \begin{cases} p \times \Delta\tau^{-1} \exp(c_2 \Delta\tau) & (\Delta\tau \lesssim 300) \\ p \times \Delta\tau^{-1} & (\Delta\tau \gtrsim 300) \end{cases} . \tag{39}$$

$c_2$ is a constant value and is roughly about $c_2 \approx 2.5 \times 10^{-2}$ (Fig E(d)).

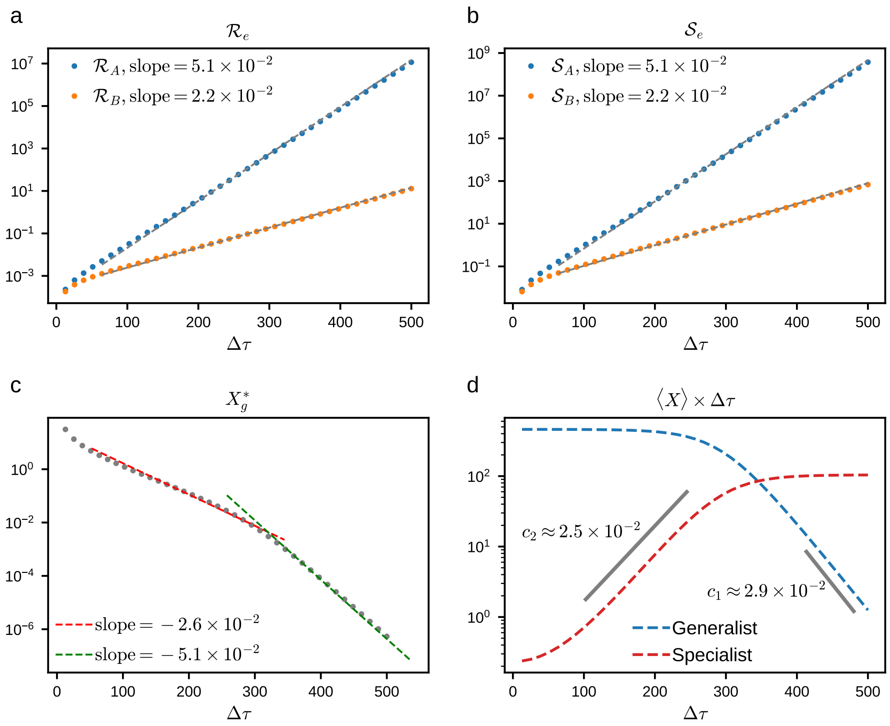


**Fig E. Order estimation.** We substituted parameter values employed in Fig.4B into analytical solutions, and plotted (a) $\mathcal{R}_e$, (b) $\mathcal{S}_e$, (c) $X_g^*$, and (d) $\langle X_1 \rangle \Delta\tau$ and $\langle X_2 \rangle \Delta\tau$ as a function of $\Delta\tau$. Assuming that each quantities follows $y = \exp(\alpha + \beta\Delta\tau)$ within a restricted range, we performed linear regression on the logarithm of each quantity to estimate the coefficient $\beta$. The gray dashed lines in (a) and (b) shows the linear regression fitted to the data points with $\Delta\tau \geq 100$. The red and green dashed lines in (c) shows the linear regression fitted to the data points with $100 \leq \Delta\tau \leq 300$ and $300 \leq \Delta\tau \leq 500$, respectively. The gray solid line in (d) shows the linear regression fitted to the data points with $100 \leq \Delta\tau \leq 300$ and $300 \leq \Delta\tau \leq 500$, respectively. All estimated values are presented to two significant figures.
